# Temporal Dynamics and Stoichiometry in Notch Signaling - from Notch Synaptic Complex Formation to NICD Nuclear Entry

**DOI:** 10.1101/2023.09.27.559780

**Authors:** Lena Tveriakhina, Gustavo Scanavachi, Emily D. Egan, Ricardo Bango Da Cunha Correia, Alexandre P. Martin, Julia M. Rogers, Jeremy S. Yodh, Jon C. Aster, Tom Kirchhausen, Stephen C. Blacklow

## Abstract

Mammalian Notch signaling occurs when binding of Delta or Jagged to Notch stimulates proteolytic release of the Notch intracellular domain (NICD), which enters the nucleus to regulate target gene expression. To determine the temporal dynamics of events associated with Notch signaling under native conditions, we fluorescently tagged Notch and Delta at their endogenous genomic loci and visualized them upon pairing of receiver (Notch) and sender (Delta) cells as a function of time after cell contact. At contact sites, Notch and Delta immediately accumulated at 1:1 stoichiometry in synapses, which resolved by 15-20 min after contact. Synapse formation preceded entrance of the Notch extracellular domain into the sender cell and accumulation of NICD in the nucleus of the receiver cell, which approached a maximum after ∼45 min and was prevented by chemical and genetic inhibitors of signaling. These findings directly link Notch-Delta synapse dynamics to NICD production with unprecedented spatiotemporal precision.

## Introduction

Notch signaling influences critical cell fate decisions in all metazoans and regulates tissue homeostasis in adults (Bray, 2016; Kovall et al., 2017; Sprinzak & Blacklow, 2021). The essential role of Notch signaling during development is evident from the embryonic lethality associated with deficiencies in Notch signaling in various model organisms, including worms, flies, and mice (Bray, 2016).

Aberrant Notch signaling is also associated with a variety of human pathologies. Germ-line mutations of core components of Notch signaling result in disorders such as Alagille syndrome, caused by loss of function mutations in *NOTCH2* or *JAGGED1* (Kamath et al., 2012; Li et al., 1997; Oda et al., 1997), and the stroke syndrome CADASIL, caused by missense mutations in the gene encoding *NOTCH3* (Joutel et al., 1996). Oncogenic gain-of-function mutations in human *NOTCH1* are frequently found in human T cell acute lymphoblastic leukemia/lymphoma (T-ALL) (Weng et al., 2004), certain B cell malignancies (Puente et al., 2011), and some solid tumors (Aster et al., 2017). Genomic studies have also uncovered loss-of-function mutations of *NOTCH1*, *NOTCH2*, and *NOTCH3* in squamous cell carcinomas in the skin, head and neck (Agrawal et al., 2011; Wang et al., 2011), and in precancerous regions of sun-exposed skin (Martincorena et al., 2015).

Mammals have four Notch receptors (NOTCH1-4) and four well-characterized activating ligands (DLL1, DLL4, JAG1, and JAG2). The Notch proteins are single-pass transmembrane receptors that are normally processed during maturation by a furin-like protease at an extracellular site called S1 (Blaumueller et al., 1997; Logeat et al., 1998) to generate non-covalently associated extracellular (NECD) and transmembrane (NTM) subunits. The mature heterodimeric receptor normally resides on the cell surface of the signal-receiving cell (or receiving cell) in an autoinhibited or “off” state and signaling is initiated at sites of cell-cell contact when Notch proteins on a receiver cell bind to Delta or Jagged ligands on a sender cell. Ligand binding relieves Notch autoinhibition by inducing proteolysis by the ADAM10 metalloprotease at a membrane proximal site called S2, producing a truncated transmembrane subunit called NEXT (for <u>N</u>otch <u>ex</u>tracellular truncation). NEXT becomes a substrate for the intramembrane protease gamma-secretase (γ-secretase), which cleaves Notch near the inner membrane leaflet at site S3. This proteolytic step releases the Notch intracellular domain (NICD), which translocates into the nucleus and forms a multiprotein complex with the DNA-binding transcription factor RBPJ, a protein of the Mastermind-like family (MAML) and additional co-activators to induce Notch target gene transcription (Bray, 2016; Sprinzak & Blacklow, 2021). The fate of the Notch extracellular domain (NECD) is less clear, but studies suggest a model in which it is endocytosed into the sender cell in complex with ligand, a process that depends on the E3 ubiquitin ligase Mindbomb (MIB) (Daskalaki et al., 2011; Guo et al., 2016; McMillan et al., 2015; Okano et al., 2016).

Although these steps of Notch signaling have been studied since *Drosophila melanogaster* Notch was cloned 40 years ago (Artavanis-Tsakonas et al., 1983), how these events are temporally coupled and choreographed during signaling is less well understood. Likewise, it is not known what the receptor-ligand stoichiometry is when complexes form at the membrane, nor is it clear how efficiently ligand-receptor engagement at the membrane leads to NICD production. Moreover, time-resolved linkage of the ligand-receptor interaction to internalization of NECD into sender cells has not been directly observed.

Using fluorescence microscopy in fly or mammalian cells transiently or stably overexpressing ligand and/or receptor molecules, others have shown that at sites of direct cell-cell contact, Notch and its ligands can gather and form stable clusters (Chapman et al., 2016; Fehon et al., 1990; Klueg & Muskavitch, 1999; Meloty-Kapella et al., 2012; Nichols et al., 2007). Similarly, transendocytosis of the NECD into vesicular structures within sending cells has also been observed in cell culture and in flies (Nichols et al., 2007; Parks et al., 2000). Ectopic overexpression of Notch can also result, however, in intracellular retention, mislocalization, and clustering of receptor molecules within the ER (Chapman et al., 2011; Mumm et al., 2000; van Tetering et al., 2009), raising the possibility that these findings are not physiologically representative. It is therefore important to use tagged Notch proteins expressed from endogenous loci to ensure faithful recapitulation of the temporal dynamics of early events responsible for Notch signaling.

In the quantitative studies reported here, we combined use of volumetric spinning disk confocal and lattice light-sheet microscopy (LLSM) (Chen et al., 2014) to image cells expressing physiological amounts of fluorescently tagged Notch and ligand proteins expressed from their endogenous loci to analyze protein localization, organization, and dynamics in living cells. LLSM was chosen because it minimizes photobleaching, increases signal to noise ratio, and allows for high spatiotemporal precision of time series recorded from the whole cell volume. When sender and receiver cells made contact, ligands and receptors clustered into synapses at the site of contact, with a synapse lifetime of roughly 15-20 min and a ligand:receptor stoichiometry of 1:1. Synapse formation preceded transendocytosis of NECD (and some full-length Notch) into the sending cell and eventual accumulation of up to 1000-2000 NICD molecules in the nucleus of the receiving cell. This work defines for the first time the stoichiometry, integrated temporal order and timing of central steps in Notch signal transduction from synapse formation through nuclear NICD accumulation and charts a course for studying real-time Notch dependent signaling dynamics in living cells in both physiological and pathophysiological contexts.

## Results

### Establishment of a system to visualize Notch signaling in real time

To study the events of physiologic Notch signaling using fluorescence microscopy in living cells, we screened for Notch- and ligand-expressing cell lines that i) were amenable to CRISPR/Cas9 engineering, ii) expressed one receptor or ligand endogenously at substantially greater natural abundance than others, and iii) were active as either receiver (Notch-expressing cells) or sender (ligand-expressing cells) cells, as assessed by assays for induction of Notch-dependent gene expression.

SVG-A immortalized fetal astrocytes met these criteria as a Notch-expressing (receiver) cell line. They have been previously successfully engineered using CRISPR/Cas9 (Chou et al., 2016), they express vastly more *NOTCH2* than other Notch isoforms (as judged by analysis of mRNA abundance by quantitative reverse-transcriptase PCR (Martin et al., 2023)), and when co-cultured with U2OS cells ectopically expressing the DLL4 ligand, they exhibit strong induction of a Notch-responsive luciferase reporter gene (Figure S1A, related to Figure 1). The reporter response was blocked by treatment with a γ-secretase inhibitor (GSI; Compound E) (Figure S1A, related to Figure 1) and was not observed in co-culture assays with parental U2OS cells. The transcriptional response to co-culture with ligand-expressing cells was also greatly reduced when *NOTCH2* was knocked out of SVG-A cells using CRISPR/Cas9 (Figure S1A, related to Figure 1), confirming that NOTCH2 was responsible for most Notch signaling activity in these cells. Importantly, when SVG-A cells were plated in tissue culture dishes containing immobilized JAG1, sentinel Notch target genes were induced within 2 to 4 hours (*e.g. HES1, TRIB1*), and the “Notch Signaling Pathway” Gene Ontology term (Chen et al., 2013; Kuleshov et al., 2016) was enriched among genes induced at 2, 4, and 24 hours after stimulation (Figure S1B, related to Figure 1).

**Figure 1.**
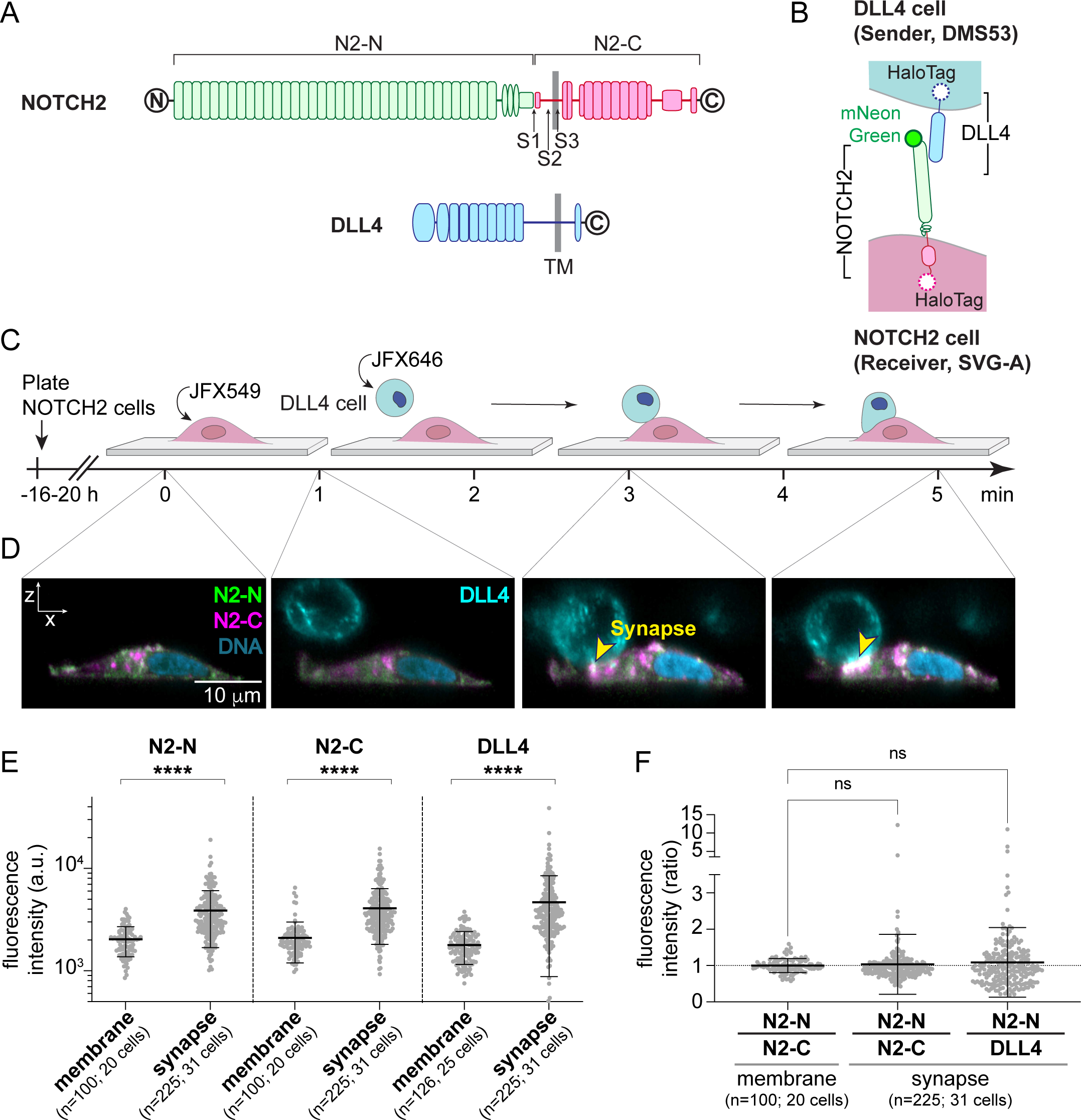
Formation of synapses at sites of NOTCH2-DLL4 contact. **A.** Domain organization of NOTCH2 and DLL4. The NOTCH2 extracellular domain (N2-N) is green, the NOTCH2 transmembrane subunit (N2-C) is magenta, and DLL4 is blue. N- and C- terminal tagging sites are shown in black. The sites of NOTCH2 proteolytic cleavage by Furin (S1), ADAM10 (S2), and γ-secretase (S3) are also indicated. **B.** Schematic showing the colors of the fluorescent labels used in cell pairing experiments. The N2-N label on NECD is mNeonGreen, the N2-C label on the C-terminal tail of NOTCH2 is HaloTag coupled to JFX549 (magenta), and the DLL4 C-terminal label is HaloTag coupled to a JFX646 (blue). **C.** Schematic of the cell paring procedure. NOTCH2 and DLL4 cells were separately labeled with JFX549 and JFX646. DLL4 cells were detached and delivered to NOTCH2 cells, and cell pairing was monitored by spinning disk confocal or lattice light-sheet microscopy. **D.** Representative lattice light-sheet images (orthogonal view, despeckle) showing NOTCH2 cells before (0 min) and 1, 3, and 5 min after microfluidic delivery of DLL4 cells. N2-N is colored green, N2-C is magenta, DLL4 is cyan, and DNA is pseudocolored blue. The synapse is indicated with a yellow arrowhead. **E.** Fluorescence intensities of N2-N, N2-C, and DLL4 signals in the regions outside of cell-cell contact (membrane) and in synapses. **F**. Ratios of fluorescence intensities of signals associated with N2-N and N2-C in the membrane and of N2-N and N2-C or N2-N and DLL4 in synapses, respectively. Data are represented as mean ± standard deviation; statistical analysis for each pair in **E** was performed using Mann-Whitney test and in **F** using Kruskal-Wallis one-way ANOVA; **** = p<0.0001, ns = p>0.05; n = number of synapses and number of cells analyzed as indicated.

We identified two ligand-expressing (signal-sending) cell lines that met our criteria. The first sender cell line was DMS53, which expresses DLL4 as its predominant ligand and is able to activate Notch as judged by the induction of reporter gene expression in SVG-A receiver cells (Figure S2, related to Figure 1). Knockout of DLL4 in DMS53 cells also reduced signal-sending activity (Figure S2H, related to Figure 1), with residual ligand activity likely resulting from the expression of other ligands (Figure S2A,B, related to Figure 1). The second sender line was A673, which endogenously expresses JAG1 as its predominant ligand (Figure S3, related to Figure 1) and induces a Notch reporter response in SVG-A receiver cells (Figure S3C, related to Figure 1). Knockout of JAG1 in A673 cells abrogated their signal sending activity (Figure S3H, related to Figure 1), consistent with the observation that JAG1 was the only ligand detectable in these cells by flow cytometry (Figure S3A,B, related to Figure 1).

**Figure 2.**
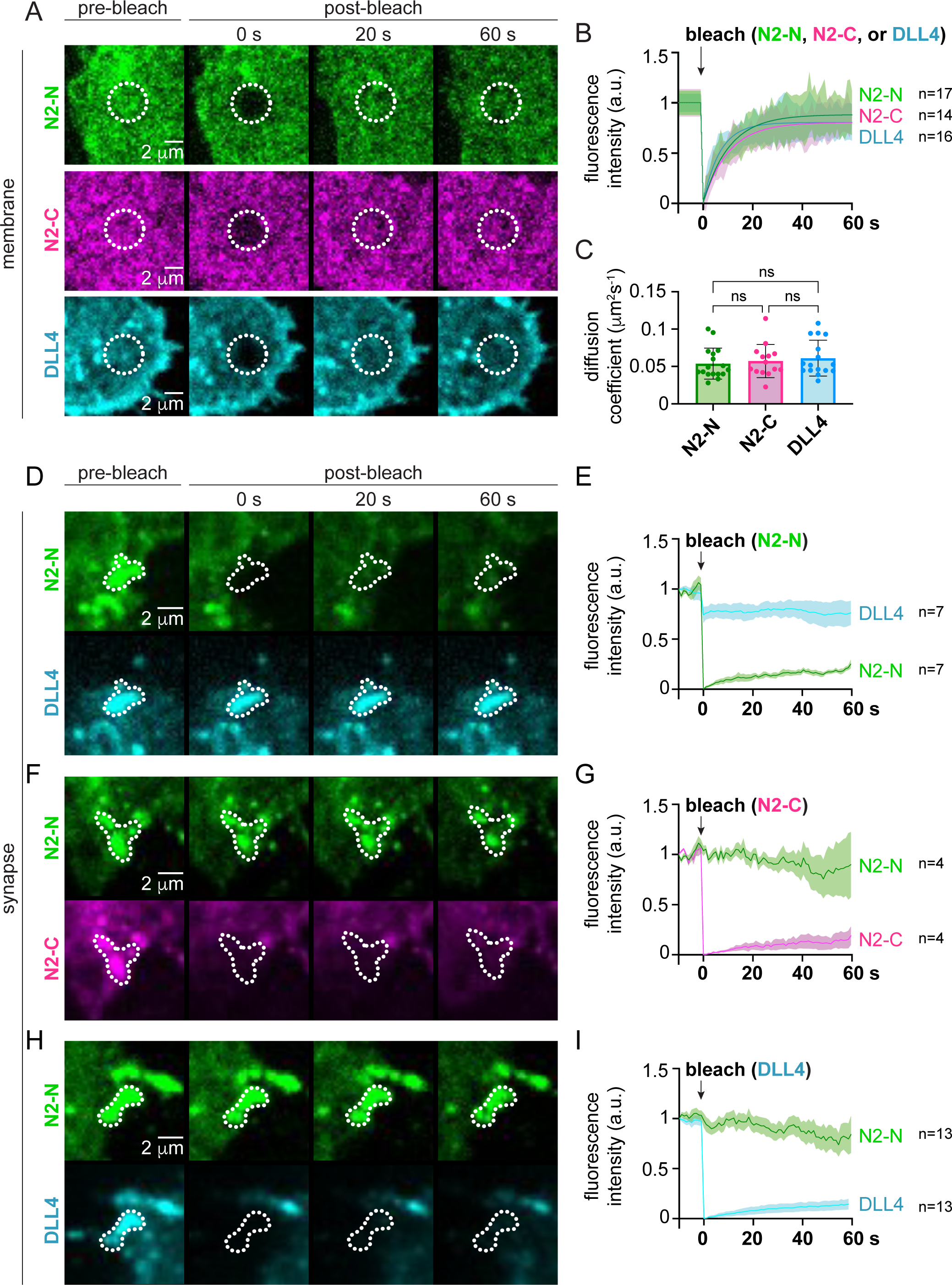
NOTCH2 and DLL4 in synapses do not readily exchange. **A.** Fluorescence recovery after photobleaching (FRAP) experiment, showing representative spinning disk confocal images of N2-N, N2-C, and DLL4 freely dispersed in the membrane before and as a function of time after photobleaching. Dotted circles indicate photobleached regions used for analysis. **B.** Recovery plots of fluorescence intensity and fitted curves (single exponential fit) after photobleaching for N2-N (green), N2-C (magenta) and DLL4 (blue) freely dispersed in the membrane. **C.** Diffusion coefficients derived from fluorescence recovery after photobleaching for N2-N (green), N2-C (magenta) and DLL4 (blue) freely dispersed in the membrane. **D, F, H.** Fluorescence recovery after photobleaching (FRAP) experiment, showing representative images of N2-N (**D**), N2-C (**F**), and DLL4 (**H**) engaged in synapses before and as a function of time after photobleaching. Images also show unbleached fluorophores (DLL4 in **D**, N2-N in **F**, and N2-N in **H**) as a positional reference for the synapses. Areas used for analysis of recovery are represented by dotted lines. **E, G, I.** Recovery plots of fluorescence intensity after photobleaching for N2-N (**E**), N2-C (**G**), and DLL4 (**I**) when engaged in synapses. Fluorescence intensity of unbleached components of the synapse (DLL4 in **E**, N2-N in **G**, and N2-N in **I**) were also monitored and analyzed as reference. Data are represented as mean ± standard deviation; statistical analysis in **C** was performed using Kruskal-Wallis ANOVA; ns = not significant; n = number of regions/synapses analyzed as indicated.

**Figure 3.**
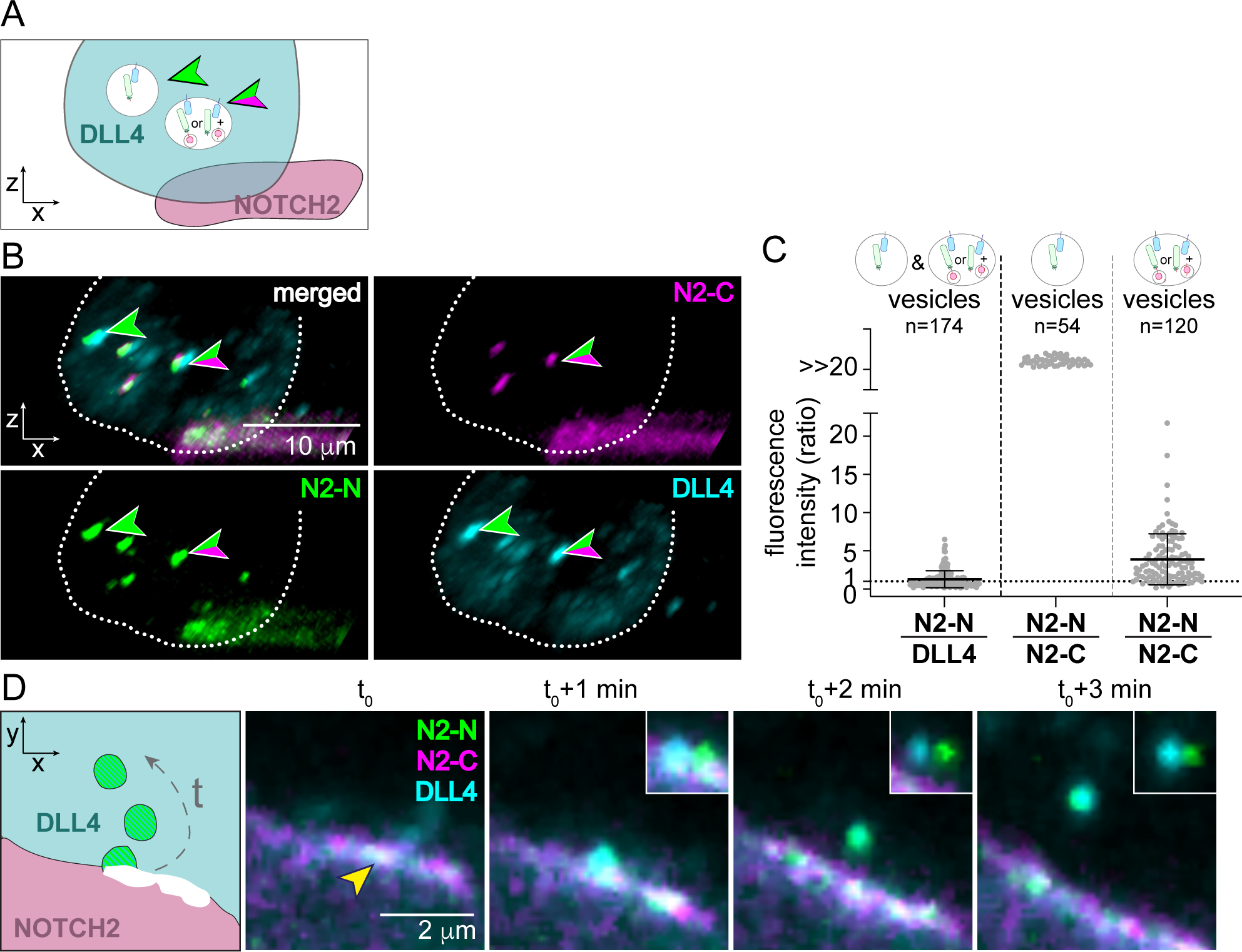
Transendocytosis of NOTCH2 into DLL4 cells takes place after synapse formation. **A.** Schematic illustrating different compositions of NOTCH2-DLL4 complexes within DLL4 cell vesicles after cell pairing. Vesicles containing N2-N:DLL4 complexes (green arrowhead) and full-length NOTCH2:DLL4 complexes (containing both N2-N and N2-C; green/magenta arrowhead) are shown. **B.** Lattice light-sheet images of a DLL4 sender cell paired with a NOTCH2 receiver cell 20 minutes after contact. N2-N in green, N2-C in magenta, and DLL4 in cyan. Green arrowhead: vesicle containing only DLL4 and N2-N fluorescence. Green/magenta arrowhead: vesicle containing DLL4, N2-N, and N2- C fluorescence. **C.** Stoichiometric ratio of N2-N to DLL4 in vesicles (left), and of N2-N/N2- C in vesicles (center and right). The stoichiometric ratio for N2-N/N2-C in vesicles where N2-C was not detectable was arbitrarily set to >>20. Dotted line indicates the ratio of one observed in membrane and synapses (see Figure 1). Number of vesicles analyzed = n. Error bars represent mean ± standard deviation. **D.** Schematic (left) and real-time lattice light-sheet microscopy images from a synapse at t0 and subsequent 1 min intervals showing movement of N2-N and DLL4 fluorescence from the synapse into the sender cell over time. Synapse at t0 is indicated with a yellow arrowhead. N2-N is shown in green, N2-C in magenta, and DLL4 in cyan. Insets show the three channels with a 5 pixel shift of the blue channel.

We used CRISPR/Cas9 in SVG-A, DMS53, and A673 cells to fuse fluorescent proteins or HaloTags (Los et al., 2008) to Notch and ligand proteins in their endogenous loci for expression at natural abundance. In SVG-A cells, NOTCH2 was double-tagged with mNeonGreen (mNeon) (Shaner et al., 2013) inserted after the signal peptide to position it extracellularly at the mature N-terminus of the NECD subunit, and with a HaloTag inserted after A2471 to place a second fluorophore intracellularly at the C-terminus of the NTM subunit (Figure 1A, Figure S4, related to Figure 1). These labeling positions are hereafter specified as N2-N and N2-C, respectively. In DMS53 and A673 cells, a HaloTag was fused to the C-terminal end of DLL4 or JAG1, respectively (Figure 1A,B, Figure S2D- F, Figure S3D-F, related to Figure 1). The steady-state expression amount and signaling activity of tagged receptor and ligand proteins were not substantially altered when compared to the endogenous proteins in parental cells, confirming that the tags do not disrupt protein processing or function (Figure S2G-J, Figure S3G-J, Figure S4E-H, related to Figure 1).

**Figure 4.**
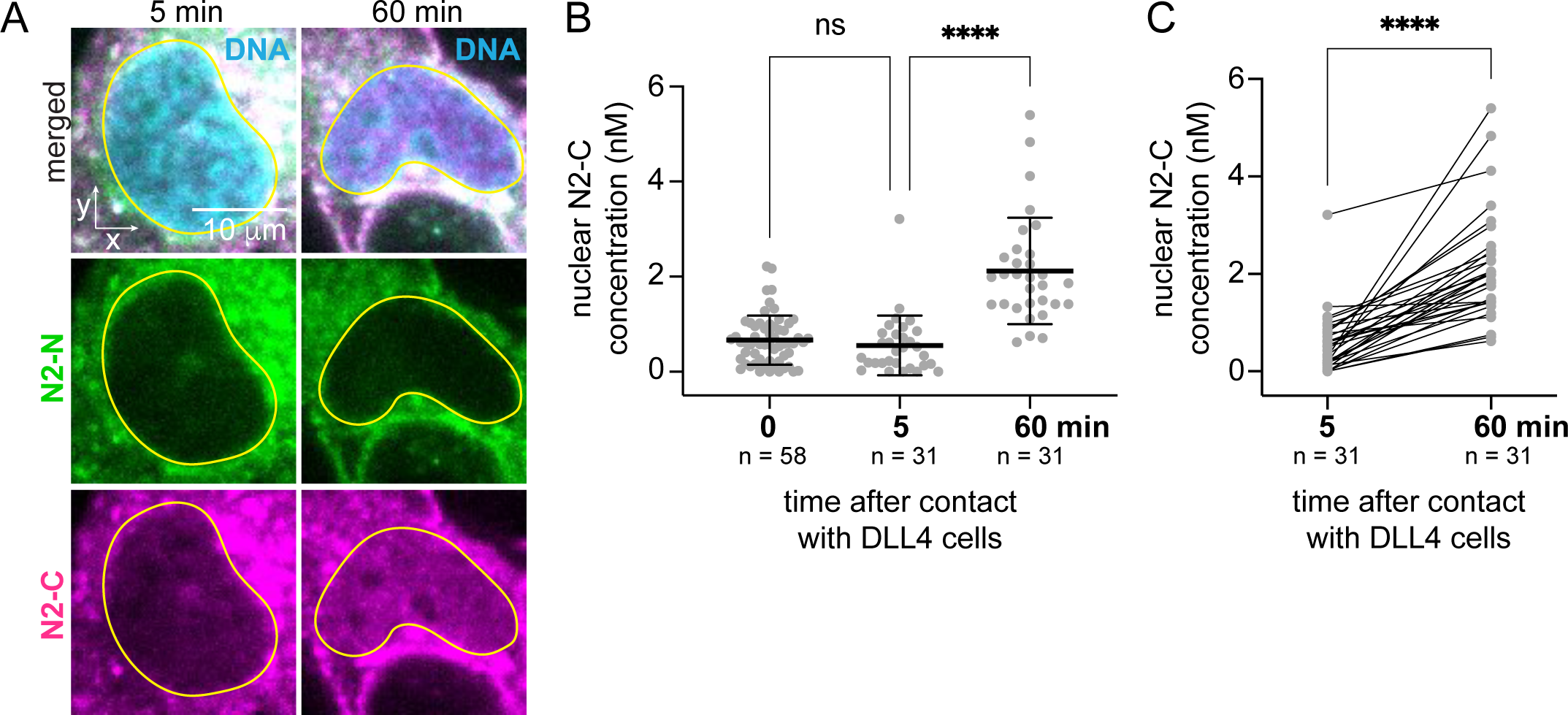
NICD nuclear entry after cell-cell contact. **A.** Representative spinning disk confocal images of a NOTCH2 cell nucleus at 5 and at 60 min after contact with a DLL4 cell. Images show the maximum intensity projection of five planes through the center of the nucleus. N2-N is green, the N2-C tag (inclusive of NTM, NEXT, and NICD species) is magenta, and the cell nucleus/DNA is pseudocolored blue (SiR-DNA). Nuclei are outlined with yellow lines. **B,C.** Quantitative analysis of the nuclear N2-C concentration (nM) before sender cell contact (0 min), and at 5 and 60 min after contact. Data are shown as a scatter plot in **B**, and lines are drawn to connect paired concentration measurements at 5 and 60 min for each nucleus analyzed in **C**. Error bars in **B** represent mean ± standard deviation. Statistical analysis in **B** was performed using Kruskal-Wallis one-way ANOVA and in **C** using Wilcoxon matched-pairs signed rank test. ns = p>0.05; **** = p<0.0001, n = number of analyzed nuclei.

We engineered a microfluidics device for imaging in a confocal (SD) or lattice light-sheet microscope (LLSM). The device made it possible to pair cells and observe the cell pairs in real time from the moment of initial contact, allowing us to follow the dynamics of NOTCH2 and DLL4 associated with signal transmission (Figure 1C, Figure S5, related to Figure 1). Sender and receiver cells were separately labeled with HaloTag ligands conjugated to different JaneliaFluorX (JFX) dyes (Grimm et al., 2020) prior to pairing. The sender cells were then delivered to receiver cells pre-plated on the cover slip by passage through a microfluidic chip using a pressure-controlled pump.

**Figure 5.**
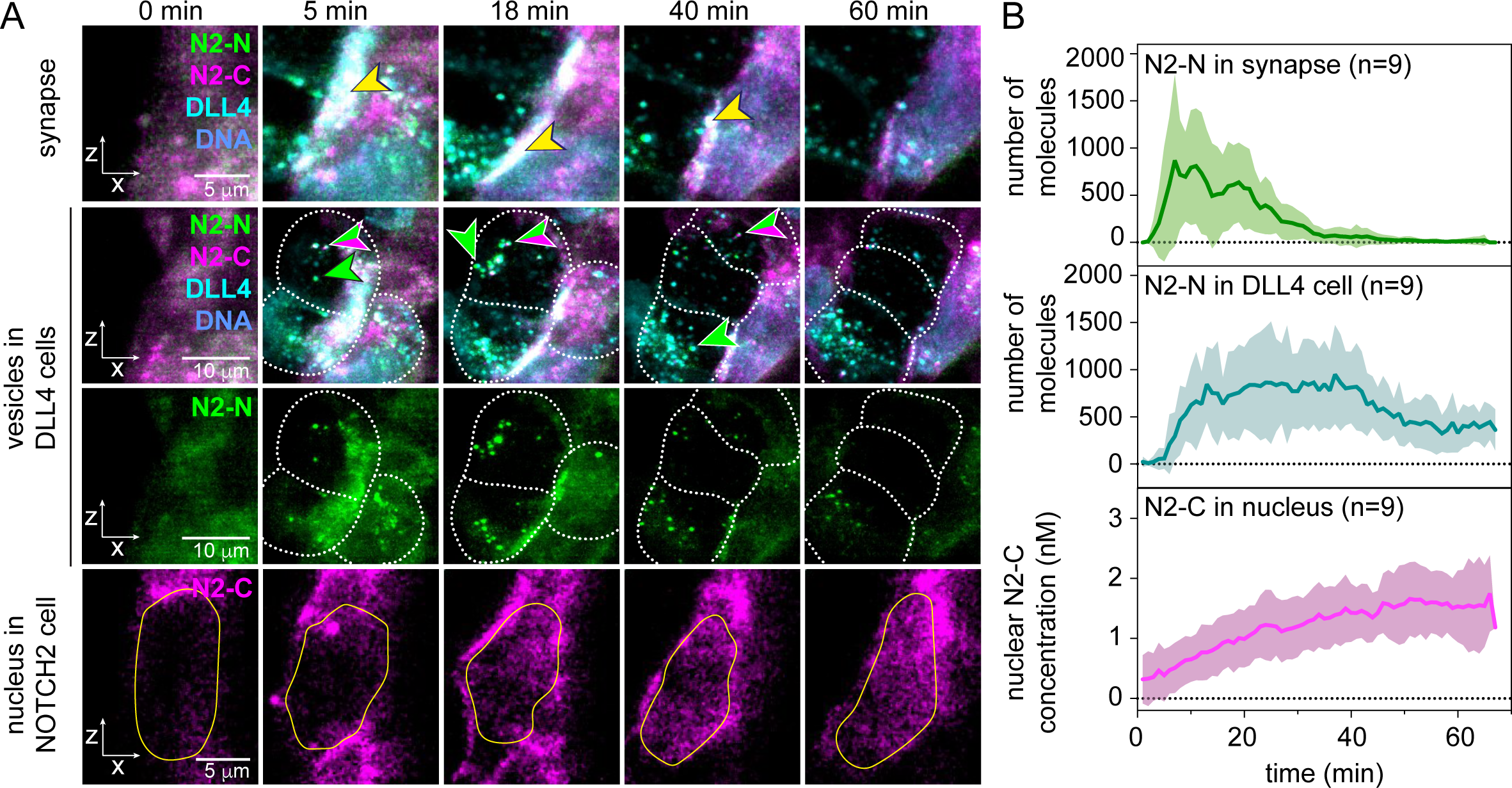
Real time visualization of events after cell pairing. **A.** Representative lattice light-sheet images from a time course observing a NOTCH2 cell before (0 min) and after contact with DLL4 cells (5-60 min). Panels highlight the formation and dissipation of synapses (top), the appearance of N2-N and N2-C positive vesicles in DLL4 cells (middle two rows) and the increase of N2-C associated signal in the nucleus of the NOTCH2 cell (bottom row). DLL4 cells are depicted by dotted lines (middle two rows) and the nucleus of the NOTCH2 cell, segmented using SiR-DNA labeling, is outlined with a yellow line (bottom row). N2-N is in green, the N2-C tag (NTM, NEXT, and/or NICD) is in magenta, and DLL4 in cyan. DNA was labeled using SiR-DNA and pseudocolored blue. Scale bars as indicated. **B.** Plots showing the estimated number of N2-N molecules in synapses (top), the number of molecules in DLL4-cell vesicles (middle) and the concentration (nM) of N2-C in nuclei of NOTCH2 cells (bottom) as a function of time after DLL4-cell contact. Graphs show mean ± standard deviation from n = 9 independent cell pairing events.

### Notch synapses form between NOTCH2 and DLL4 at sites of cell-cell contact

In cultured SVG-A cells, NOTCH2 was found at the plasma membrane and in intracellular puncta (Figure 1D, Figure S6A, related to Figure 1) that likely represent trafficking vesicles and/or organelles related to protein synthesis and degradation. The concurrent presence of nonspecific or autofluorescence signals in the green (488) and red (561) channels, also seen as small intracellular puncta in both parental cells and in knockin cells that did not have a JFX dye coupled to the HaloTag (Figure S6A,B, related to Figure 1), prevented unambiguous identification of NOTCH2-containing vesicles inside these cells.

DMS53 sender cells delivered to the SVG-A receiver cells allowed real time imaging of DLL4 engagement with NOTCH2 at sites of contact (Figure 1C,D). These sites, which we defined as Notch synapses, showed accumulation of NOTCH2 and DLL4 and presumably occurred at sites of molecular contact between the ectodomains of DLL4 and NOTCH2 (Figure 1D, Movies 1, 2A,B). Synapses formed with 100% efficiency within seconds every time these two cell types made direct contact and varied in size and shape (Figure S7A, related to Figure 1). Preincubation of DMS53 cells with ligand-blocking antibodies prevented synapse formation and effectively silenced signaling (Figure S7B,C, related to Figure 1), indicating that synapse formation required direct binding of DLL4 to NOTCH2.

To evaluate whether the proteins concentrated at points of cell-cell contact, we compared the fluorescence intensities of the N2-N, N2-C, and DLL4 tags in synapses to their intensities in membrane regions excluded from the synapses (“membrane”) and measured significantly higher fluorescence intensity signals in the synapses (Figure 1E).

We determined the ratio of fluorescence intensities of the N2-N and N2-C tags in the membrane of receiving cells (before delivery of ligand cells), and set the value of that ratio to a stoichiometry of 1:1 because both fluorophores are coupled to the same receptor protein. The same 1:1 stoichiometry was observed outside synapses after Notch cells contacted sender cells (Figure 1F; N2-N/N2-C in membrane). The N2-N:N2-C stoichiometry remained 1:1 in synapses associated with NOTCH2 - DLL4 engagement (Figure 1F; N2-N/N2-C in synapse). To determine the stoichiometric ratio of NOTCH2 to DLL4 in synapses, we exploited the capacity of the HaloTag to be labeled with different dyes and exchanged the Notch C-terminal and DLL4 fluorophores to determine the NOTCH2:DLL4 ratio in the synapse. We established that the N2-N to N2-C and N2-N to DLL4 fluorescent tag ratios were 1:1 independent of the dyes exchanged and indistinguishable from each other (Figure 1F; N2-N/N2-C and N2-N/DLL4 in synapse).

Similarly, a 1:1 receptor:ligand stoichiometry was present at synapses formed by NOTCH2 and JAG1 upon pairing A673 JAG1-HaloTag cells and NOTCH2-tagged SVG- A cells (Figure S3L,M, related to Figure 1). One detectable difference was that the A673 (JAG1) cells formed synapses less efficiently than the DMS53 (DLL4) cells (Figure S3N, related to Figure 1), most likely because the amount of JAG1 on the surface of A673 cells was lower than the amount of DLL4 on DMS53 cells. In each case, endogenously expressed ligands and receptors formed synapses at contact sites in living cells with a stoichiometry of 1:1.

### NOTCH2 and DLL4 in synapses do not readily exchange

We performed fluorescence recovery after photobleaching (FRAP) in a spinning disk confocal microscope to assess the dynamics of receptor and ligand exchange on the cell surface, both in regions outside of and within synapses. FRAP was performed within a region of interest (ROI) and recovery was monitored at 1 s intervals for a total of 60 s. Outside sites of cell contact, the fluorescence intensity after bleaching recovered 80%- 90% of the initial value after 60 s for both the N2-N and N2-C tags and for the DLL4 tag, indicating that both proteins are mobile on the cell surface (Figure 2A,B). The half-times for recovery (t1/2) of N2-N and N2-C on SVG-A cells were 7.5±2.5 and 7.2±3.0 s, which correspond to diffusion coefficients (D) of 0.053±0.02 μm^2^s^-1^ and 0.057±0.02 μm^2^s^-1^, respectively (Figure 2B,C). Free DLL4 molecules on the surface of DMS53 cells had a similar mobility, with a recovery t1/2 of 4.7±1.6 s and a diffusion coefficient of 0.061±0.024 μm^2^s^-1^ (Figure 2B,C). These diffusion coefficients are comparable to that of stably overexpressed DLL1 in CHO-K1 cells (Khait et al., 2016) and to those of other freely diffusing membrane proteins (Jacobson et al., 1987).

We next determined the mobility of Notch and DLL4 molecules at the synapse by bleaching the fluorophore of interest 5-10 min after the onset of synapse formation. We monitored the fluorescence intensity of the non-bleached component within the region of interest (ROI) to delineate the synapse’s location and ascertain its structural integrity throughout the 60-second observation period (Figure 2D,F,H). In contrast to the rapid fluorescence recovery of N2-N, N2-C or DLL4 in the surrounding cell surface membrane, NOTCH2 or DLL4 did not readily exchange when in synapses (10-20% recovery after 60 s) (Figure 2E,G,I). Thus, at the site of contact, both receptor and ligand exhibited greatly reduced exchange within the synapse and/or with the surrounding membrane.

### Notch transendocytosis into the sender cell occurs after synapse formation

Transendocytosis of NECD and full-length Notch into ligand cells has been observed in cultured cells (Nichols et al., 2007; Parks et al., 2000) and in flies (Langridge & Struhl, 2017; Parks et al., 2000). Here, we monitored transendocytosis of NOTCH2, which accumulated into puncta within DMS53 (DLL4) cells only after synapse formation between paired cells (Figure 3A,B, Movie 3). We quantified the relative amounts and stoichiometry of the N2-N (*i.e.* NECD) and N2-C tags to the DLL4 tag in these puncta by determining the fluorescence intensity ratios of different fluorophore pairs in these structures 60 min after synapse formation. The N2-N (*i.e.* NECD) to DLL4 stoichiometric ratio was approximately 1:1 (Figure 3C; left, n = 174 puncta) in agreement with the ratio of receptor to ligand in synapses. In 54 of the puncta, only the N2-N (*i.e.* NECD) and DLL4 were detected (Figure 3C; middle), whereas in the other puncta some N2-C was present along with N2-N and DLL4 (Figure 3C; right), indicative of occasional transendocytosis of full-length NOTCH2 as well as just the NECD. Quantification of the N2-N/N2-C ratio in these puncta showed an average value of 4:1, with considerable variation among the puncta.

While the majority of transendocytosis events involved only or predominantly N2-N (*i.e.* NECD), the entry of some N2-C into ligand cells along with N2-N suggested that some non-productive transendocytosis of full-length receptors occurred. Consistent with this interpretation, we did not observe any evidence of Notch signaling activity when ligand (DMS53) cells were probed using a luciferase reporter for a NICD-dependent response (Figure S8, related to Figure 3), and we did not observe any accumulation of NICD in the nuclei of those cells. We also did not detect entry of DLL4 into the SVG-A (NOTCH2) receiver cells.

Occasionally, we were able to observe vesicle-like structures containing both ligand and receptor adjacent to synapse sites (Figure 3D). While our analyses did not allow us to determine unambiguously whether these vesicles originated directly from synapses or if the NOTCH2 and DLL4 instead accumulated in vesicles residing close to the contact site, it is possible these objects are vesicles captured at a very early stage shortly after initiation of transendocytosis.

### Quantification of nuclear entry of NICD after cell contact

NICD can access the nucleus within 30 min of γ-secretase inhibitor removal (Martin et al., 2023) and can induce a transcriptional response in the nucleus within 60 min of Notch activation (Falo-Sanjuan & Bray, 2022; Ilagan et al., 2011). To quantify the amount of N2- C entering nuclei after cell-cell contact, we paired and imaged sender and receiver cells immediately (1-5 min) and 60 min after cell contact. Visual inspection of the nuclear region showed an increase of the fluorescent signal of N2-C, consistent with NICD nuclear entry (Figure 4A). The nuclear N2-C (*i.e.* NICD) concentration, calculated using a calibration curve with purified, recombinant HaloTag protein in solution labeled with JFX549 (Figure S9, related to Figure 4), rose from 0.65±0.6 nM before or immediately after cell contact to ∼2±1.1 nM (equivalent to ∼1000-2000 NICD molecules) at a time point 60 minutes after synapse formation (Figure 4B,C). The presence of intracellular puncta in the isolated NOTCH2 cells did not allow us to unambiguously follow the path of N2-C (*i.e.* NICD) from the synapse to the nucleus.

### Temporal linkage between Notch processing and nuclear entry in living cells

We next established a quantitative spatiotemporal link among synapse formation, NECD transendocytosis, and NICD nuclear accumulation by using our microfluidic device to obtain imaging data of nine cell pairing events with a lattice light-sheet microscope over a 60-minute time course (Figure 5 and Figure S10, related to Figure 5). This approach enabled three-dimensional (3D) visualization with little photobleaching and phototoxicity compared to conventional spinning disk microscopes, thereby allowing repeated quantitative imaging of fluorescently tagged proteins expressed at endogenous levels over a prolonged period of time. The signal distribution of NOTCH2 at the cell surface was homogeneous in the absence of contact with DMS53 sender cells (t=0), as assessed by analysis of N2-N and N2-C tag fluorescence, and the nuclear N2-C signal was minimal (Figure 5A,B). Again, Notch synapses rapidly formed at the site of contact between sender and receiver cells; NOTCH2 and DLL4 molecules accumulated within seconds after contact and the average synapse grew (assessed by the N2-N signal) from roughly 500 NOTCH2 molecules after 5 min of contact to a peak of roughly 2000 molecules at 15-20 min. After 30 min, the synapses typically resolved (Figure 5 and Figure S10, related to Figure 5). The number of N2-N (*i.e.* NECD) molecules in puncta of DMS53 sender cells increased to a maximum at roughly 15 minutes before slowly decaying after 40 min, perhaps due to protein degradation, entry into a compartment with a different pH, or both (Figure 5). Finally, the concentration of N2-C (*i.e.* NICD) in the nuclei of the receiver cells increased to a maximum of 1.42±0.41 nM, corresponding to 1000-2000 molecules ∼45 minutes after cell-cell contact, and remained steady until the end of the 60 min time course (Figure 5).

### Mindbomb, ADAM10, and γ-secretase are not essential for synapse formation but are required for nuclear entry of NICD

The E3 ubiquitin ligase Mindbomb1 (MIB1) is required in sender cells for ligand activity and subsequent receptor activation (Guo et al., 2016). We eliminated *MIB1* in DLL4- HaloTag cells (*MIB1ko*) using CRISPR/Cas9 (Figure S11A,B, related to Figure 6) and paired these cells with our tagged SVG-A cells to monitor synapse formation, Notch transendocytosis, and N2-C accumulation in the nuclei of Notch cells. *MIB1ko* cells formed synapses efficiently but these synapses did not resolve after 60 min (Figure 6A). *MIB1ko* cells were also unable to induce transendocytosis of N2-N (*i.e.* NECD) (Figure 5B), and failed to produce a substantial increase in nuclear N2-C (*i.e.* NICD) within receiver cells (Figure 6C,D). These data show that MIB1 in sender cells is essential for synapse dissolution, and confirm it is required both for endocytosis of ligand-NECD complexes into the sender cell and for nuclear entry of NICD in the receiver cell.

**Figure 6.**
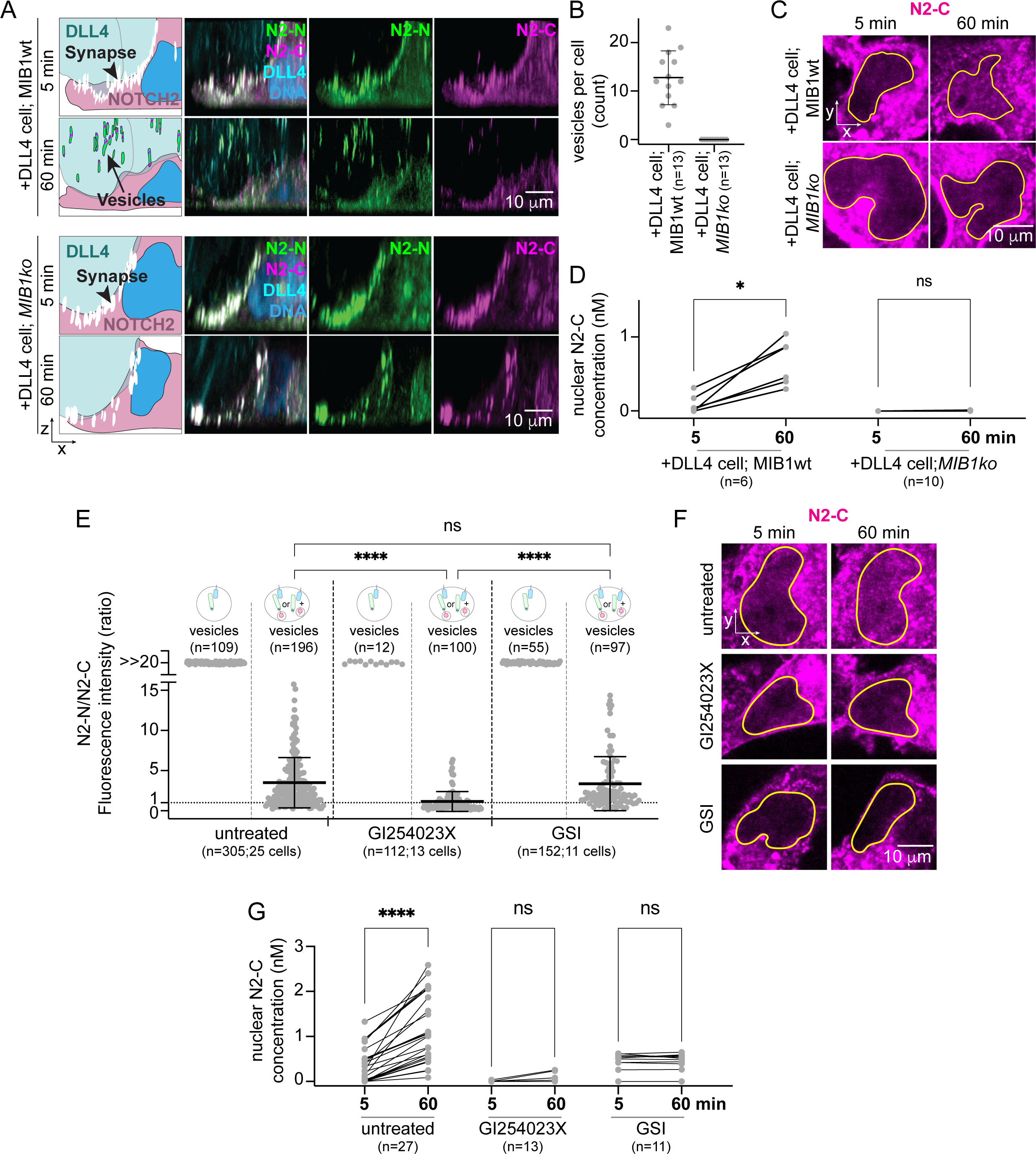
Effects of chemical and genetic perturbations on synapse formation, transendocytosis and nuclear NICD entry. **A-D.** Effects of knocking out MIB1 (*MIB1ko*) in sender cells. **A.** Pairing of parental (top) and *MIB1ko* (bottom) DLL4 sender cells with NOTCH2 receiver cells, imaged using a spinning disk confocal microscope. Schematics (left) show cells, synapses (white, indicated by the black arrowhead), vesicles in DLL4 cells (black arrow), and nuclei (blue) of NOTCH2 cells. Images (right) show paired cells 5 and 60 min after contact. N2-N is shown in green, N2-C in magenta, DLL4 in cyan, and the nucleus of the NOTCH2 cell is pseudocolored blue. Images show the maximal intensity projection of a 3D z-stack of 14.84 μm. **B.** Vesicles per DLL4 cell (MIB1 parental or *MIB1ko*) 60 min after NOTCH2 cell contact, assessed by manual counting. **C.** Representative images of nuclei from NOTCH2 cells co-cultured with parental or *MIB1ko* DLL4 sender cells, shown 5 and 60 min after direct contact. N2-C is shown in magenta. The images show the maximum intensity projection of five planes through the center of the nucleus. Yellow outlines denote nuclei as segmented using SiR-DNA labeling. **D.** Quantitative analysis of the N2-C concentration (nM) in nuclei from NOTCH2 cells co-cultured with parental or *MIB1ko* DLL4 sender cells at 5 and 60 min after direct contact.s**E.** N2-N/N2-C stoichiometric ratios in DLL4-containing vesicles of sender cells co-cultured with untreated, GI254023X-treated, or GSI-treated NOTCH2 cells. **F.** Representative images of nuclei from untreated, GI254023X-treated, or GSI-treated NOTCH2 cells at 5 and 60 min after direct contact with DLL4 cells. N2-C is shown in magenta. Yellow outlines denote nuclei as segmented using SiR-DNA labeling. Each image shows the maximum intensity projection of three planes through the center of the nucleus. **G.** Quantitative analysis of the N2-C concentration (nM) in nuclei of untreated, GI254023X-treated or GSI- treated NOTCH2 cells at 5 and 60 min after contact with sender cells. Error bars in **B** and **E** show mean ± standard deviation. Statistical analyses in **D** and **G** were performed using the Wilcoxon matched-pairs signed rank test, and in **E** using Kruskal-Wallis one-way ANOVA. Dotted line in **E** indicates the ratio of one observed in membrane and synapses (Figure 1). ns = p>0.05; ** = p<0.001, **** = p<0.0001, n = number of analyzed cells, nuclei or vesicles as indicated.

We used protease inhibitors to investigate how preventing ADAM10 or γ-secretase cleavage of Notch affects the behavior of DLL4 and NOTCH2 after cell pairing. While synapses still rapidly formed after contact (Figure S11C, related to Figure 6), they resolved when cleavage at S2 was prevented with the metalloprotease inhibitor GI254023X. This resolution may be due in part to transendocytosis of intact NOTCH2 into the sender cells because the ratio of signals from the N2-N and N2-C labels was 1:1 in the internalized structures (Figure 6E), indicating that ADAM10 inhibition did not interfere with the transendocytosis of full-length receptors. As expected, accumulation of N2-C (*i.e.* NICD) in receiver cell nuclei was greatly reduced (Figure 6F,G). Sender and receiver cell pairs also formed Notch synapses that resolved within 60 min in the presence of a γ- secretase inhibitor (GSI; Compound E). Under these conditions, transendocytosis of N2- N (*i.e.* NECD) and full-length NOTCH2 into sender cells was not affected when compared to untreated cells (Figure 6E), indicating that release of the NECD by ADAM10 proteolysis was still occurring. As expected, we failed to observe any increase in the nuclear content of N2-C (*i.e.* NICD) even 60 min after initiation of cell-cell contact (Figure 6F,G); these observations confirmed that γ-secretase was required for the cleavage step that produces NICD and for its subsequent entry into the nucleus.

## Discussion

In this work, we directly visualized NOTCH2 and DLL4 proteins from the onset of contact between DLL4 sender and NOTCH2 receiver cells until nuclear NICD in the receiver cells accumulated to steady state. A critical feature of this study was the use of genome edited cells to ensure that the fluorescently tagged proteins were present at their natural abundance. Using quantitative fluorescence microscopy, we uncovered the appearance of a transient structure at the contact site between DLL4 sender and NOTCH2 receiver cells, here termed a Notch synapse. The Notch in the synapse is the source of the NICD that accumulates in the nucleus of the receiver cell.

Notch synapses form immediately after signal sending and signal receiving cells meet, as previously observed at contact sites in other model systems that used ectopic protein overexpression (Chapman et al., 2016; Fehon et al., 1990; Khamaisi et al., 2022; Meloty-Kapella et al., 2012; Nichols et al., 2007). In contrast to our work, which uncovered the transient presence of a Notch synapse elicited immediately after sender-receiver cell contact, the previous studies using overexpressed proteins instead observed stable synapses that could last 24 hours or longer after their formation (Chapman et al., 2016; Fehon et al., 1990; Khamaisi et al., 2022).

Strikingly, NOTCH2-DLL4 synapses accumulated normally but failed to resolve in synapses created between Notch receiver and sender cells lacking the E3 ligase MIB1. Because NECD (represented by the N2-N tag) from SVG-A sender cells failed to transendocytose into DMS53 (DLL4) *MIB1ko* cells, we disfavor a previous model for activation in which the furin-processed extracellular and transmembrane subunits of Notch are mechanically induced to dissociate at site S1 prior to metalloprotease cleavage (Chastagner et al., 2017; Nichols et al., 2007). Additionally, a model in which mechanical force supplied by bound ligand induces subunit dissociation at site S1 also predicts that ADAM10 inhibition would still be permissive of transendocytosis of liberated NECD into the sender cells, yet we observed that - although treatment with an ADAM10 inhibitor allowed transendocytosis of full-length NOTCH2 into the sender cells - it failed to permit transendocytosis of the free NECD. Our data are instead consistent with models positing that MIB1-dependent endocytosis of ligand is needed to induce ADAM10 cleavage of Notch at S2 in receiver cells, thereby liberating NECD. Our results are also consistent with findings in flies, in which replacement of the Notch negative regulatory region (NRR), which contains the S1 and S2 cut sites, by a domain more resistant to force-induced unfolding also leads to transendocytosis of full-length receptors, but not free NECD, into sender cells (Langridge & Struhl, 2017). Unlike the studies in flies, however, in which ligands could enter the cells expressing the unfolding-resistant chimeric receptors, we did not observe entry of any DLL4 into the SVG-A receiver NOTCH2 cells when ADAM10 cleavage was chemically inhibited.

We showed that both NOTCH2 and DLL4 were mobile when on the cell surface but became fixed at the contact site once synapses formed. The mobility of DLL4 and NOTCH2 outside sites of contact resembled that predicted for their lateral diffusion in the membrane, and was similar to that of overexpressed DLL1 in the membranes of CHO cells (Khait et al., 2016). The relative immobility of the molecules in synapses suggests the existence of avidity effects that hold the molecules in place at the observed 1:1 stoichiometry (Figure 1).

Whether the stabilization of molecules in the synapses is a consequence of structured polymerization or another mechanism of self-association among the NOTCH2 and DLL4 molecules is not clear. There is evidence for weak self-association of the ankyrin domains of *Drosophila* Notch (Allgood & Barrick, 2011) and human NOTCH1, which contribute to the cooperative formation of dimeric transcription complexes on paired site DNA (Arnett et al., 2010). It is also true that the negative regulatory regions (NRRs) from NOTCH1, NOTCH2, and NOTCH3 share a crystal packing interface, the disruption of which induces signaling independent of ligand-receptor interaction (Gordon et al., 2009, 2007; Xu et al., 2015). However, the surface density of NOTCH2 and DLL4 in synapses appears to have been too low for them to be the only proteins present in synaptic sites, suggesting that additional proteins are needed to form the scaffold that holds them in a synapse.

NICD accumulation could be observed in the nucleus of receiver cells as early as 10 min after contact and plateaued after roughly 45 min. Live imaging of GFP-tagged Notch in sensory organ precursor cells of flies has shown that Notch can be seen in the nucleus on the pIIa cell as early as 10 min after cell division of the pIIa/pIIb precursor (Couturier et al., 2012). The accumulation of steady-state levels of NICD in the nucleus by ∼45 min is also in agreement with the observed timing for transcriptional induction of Notch target genes in *Drosophila* and cell culture systems (Falo-Sanjuan et al., 2019; Ilagan et al., 2011; Pillidge & Bray, 2019). The timing of these dynamics are also similar to that obtained by following the kinetics of proximity labeling of nuclear proteins associated with the Notch transcriptional response which become labeled within 30-45 min of release from GSI inhibition (Martin et al., 2023).

NICD entry into the nucleus of the receiver cell only occurred after synapse formation and only when NECD entry into the sender cell was also observed. The relatively uniform nuclear distribution of NICD, outside of nucleoli, from which it appeared to be excluded, made it possible to estimate the number and concentration of NICD molecules in the nucleus. Because the distribution of NICD in the receiver cell nuclei was not punctate, it appears that NICD does not accumulate in transcriptional hubs or nuclear foci, and that formation of such foci are thus not required for transcriptional induction in response to NICD, at least in the first hour after cell contact.

More broadly, our studies illustrate the power of real-time imaging associated with signaling dynamics using proteins labeled at natural abundance. Using this approach, we uncovered dynamic formation and dissolution of synapses at sites of cell contact, quantified the stoichiometry of ligand-receptor complexes in synapses, and saw directly that synapse formation preceded transendocytosis of NECD into the sender cell, followed by entry of NICD into the nucleus of the receiver cell. Application of this strategy to other signal transduction systems should facilitate deeper understanding of their dynamics and molecular mechanisms with potential to make important new contributions in the analysis of complex biological systems during cell differentiation *in vitro* and *in vivo* with unprecedented spatiotemporal precision.

## Acknowledgments

We thank all members of the Blacklow and Kirchhausen laboratories for helpful discussions and encouragement. We thank Matthieu Delincé for design of the PDMS chip architecture, Luke D. Lavis and Jonathan B. Grimm for the Janelia Fluor HaloTag ligands, Tegy John Vadakkan and Eric Marino for maintaining the spinning disk confocal microscope and the Center for Nanoscale Systems at Harvard University where we produced the microfluidics devices. This work was supported by NIH awards 1R35 CA220340 (to S.C.B.), R01 CA272484 (to S.C.B. and T.K.), 5R35 GM130386 (to T.K.), and K99 GM144750 (to J.M.R.). L.T was supported by a DFG Walter-Benjamin fellowship (TV-11/1). J.C.A. was supported by the Ludwig Center at Harvard.

## Author contributions

L.T., G.S., T.K and S.C.B. conceived the project. S.C.B. and T.K. acquired funding. L.T., G.S., E.D.E., R.B.D.C.C. performed experiments. L.T and G.S. analyzed the data. E.D.E. and J.M.R. processed and analyzed RNA-Seq data. A.P.M. generated A673 *JAG1ko* cells, J.S.Y. helped with the fabrication of microfluidic chips, A.P.M and J.C.A. assisted with data analysis and interpretation. L.T., G.S., T.K. and S.C.B. wrote the manuscript with input from all authors. All authors provided feedback and agreed on the final manuscript.

## Materials and Methods

### Cell culture

All cell lines were cultured at 37°C and 5% CO2 in DMEM supplemented with 10% heat inactivated fetal bovine serum (FBS, GeminiBio, 100-106) and 100 U/ml penicillin and streptomycin (ThermoFisher Scientific, 15140163) unless otherwise specified. All cell lines were periodically tested for mycoplasma by PCR. Cells were detached from plates after a PBS rinse using 0.05% Trypsin/0.53 mM EDTA in HBSS (Corning) for 5-10 min at 37°C unless otherwise specified.

### Genome editing

CRISPR/Cas9 was used for genome editing to engineer doubly tagged NOTCH2 in SVG-A cells. mNeonGreen flanked by GGS (gly-gly-ser) linkers was inserted after the signal peptide of NECD; HaloTag was inserted at the C-terminus of NTM after a GGAG (gly-gly-ala-gly) linker sequence and immediately before the stop codon. CRISPR/Cas9 editing was also used to insert a HaloTag at the C-terminus of DLL4 in DMS53 cells and at the C-terminus of JAG1 in A673 cells. Halo Tag was placed between a GGAG linker and immediately before the stop codon in A673 cells, or between a GGAG linker and a T2A sequence preceding a neomycin resistance cassette in DMS53 cells. Parental cell lines were seeded onto 6-well plates and transfected the next day with a mixture of repair template (8 µg) and a pX459 plasmid (4 µg) encoding the single guide RNA (gRNA) and *S. pyrogenes* Cas9 using Lipofectamine™ 2000 (Invitrogen). Single SVG-A or A673 cells were sorted by fluorescence (mNeonGreen or HaloTag labeled with JFX646) using a Sony SH800S Cell Sorter (Sony Biotechnology) six days after transfection and collected in 50:50 conditioned:complete media (SVG-A) or 50:50 conditioned media:FBS (A673). Single colonies of DMS53 cells were obtained by selection for 30 days using DMEM supplemented with 15% FBS (Gibco, 10437028), 100 U/ml penicillin and streptomycin and G418 (1 mg/ml; Geneticin, Gibco). Colonies were manually picked and expanded. Successful tag integration in single colonies of all cell lines was detected using genome-specific primers and PCR-based genotyping. The correct sequence was then confirmed by Sanger DNA sequencing of the PCR-amplified region.

Knockout of *NOTCH2* in SVG-A cells was performed by gRNA targeting of the sequence downstream of the signal peptide in exon 2, and knockout of *JAG1* in A673 cells was carried out with two gRNAs flanking exon 1. The gRNAs were subcloned into pX458, which contains an eGFP coding sequence behind a T2A cassette downstream of the gRNA insert. SVG-A or A673 cells were transfected with the gRNA-containing plasmids using Lipofectamine™ 2000 (Invitrogen), and cells were allowed to grow for 3-6 days. Cells were then sorted for eGFP fluorescence (indicative of plasmid uptake) using a SONY SH800S Cell Sorter (Sony Biotechnology). Single SVG-A or A673 green cells were collected in 50:50 conditioned:complete media or 50:50 conditioned media:FBS, respectively. Cells were expanded and gene editing was confirmed by genotyping and Western Blot analyses. For knockout of *DLL4* or *MIB1* in DMS53 cells, two sgRNAs flanking exon1 of the target gene were subcloned into pX459 plasmids containing a puromycin resistance (puroR) gene. Cells were transfected with plasmids carrying the sgRNAs and were incubated in DMEM supplemented with 15% FBS (Gibco, 10437028), 100 U/ml penicillin and streptomycin, and puromycin (10 ug/ml) for 3 days. Puromycin was removed and single colonies were allowed to grow for 30 days. Subsequently, colonies were manually picked, expanded, and screened for DLL4 or MIB1 loss using anti-DLL4 or anti-MIB1 antibodies by Western blot and for DLL4, by flow cytometry.

### JAGGED1-Fc expression and purification

Human JAGGED1-Fc (Martin et al., 2023) was transfected into Expi293F cells (ThermoFisher, A14527) using FectroPro (Polyplus, 101000007). Secreted JAGGED1-Fc was recovered from the culture media on Protein A agarose (Millipore, 16-125) and eluted with 100 mM glycine, pH 3.0. The eluate was neutralized with 1M HEPES buffer pH 7.3, concentrated, and buffer exchanged into 20 mM HEPES pH 7.3, containing 150 mM NaCl and 10% glycerol.

### RNA seq sample preparation

SVG-A cells were removed from plates by treating with 0.5 mM EDTA for 3 min, quenched with media, and counted. 4 x 10^5^ cells per well were plated in media containing 100 nM GSI (Compound E; Millipore, 565790) on non-tissue-culture treated 6-well plates that were pre-treated overnight with PBS + 0.1 mg/ml Poly-D-lysine (Thermo Scientific, A3890401) and 200 μg/ml human JAGGED1-Fc. After 18 h, the SVG- A cells were washed three times in 4 ml of media to remove GSI, and incubated for 2, 4, or 24 hours before harvesting by resuspension in 1 ml Trizol (Thermo Scientific, 15-596- 026). A “0 hr” reference control was collected by performing a mock washout with media containing 100 nM GSI and immediately harvesting in Trizol.

### RNA seq library construction

Samples in Trizol were thawed, and ERCC spike-in RNAs (Thermo Scientific, 4456740) were added at 10 μl per million cells. RNA was isolated using chloroform following the MaXtract tube protocol (Qiagen 129056). 5 μg of RNA was treated with DNaseI (Thermo Scientific, 18068015) in the presence of SUPERase-In (Thermo Scientific, AM2696). RNA quality was evaluated by HS RNA ScreenTape (Agilent, 5067-5579) on a TapeStation; all samples had RIN score > 8. 500 ng RNA was used as input for the TruSeq Stranded Total RNA sequencing kit with RiboZero rRNA depletion (Illumina, 20020598). Samples were sequenced at the Harvard University Bauer Core on a NovaSeq 6000 using the S1 300 cycle kit, with paired end 150 bp reads.

### RNA seq analysis

Reads were first mapped to ERCC spike in sequences using bowtie1.2.2 with the following parameters: -n2 -l 40 -X1000 --best -3 (Langmead et al., 2009). Reads not mapping to the spike-in sequences were mapped to hg38 using STAR version 2.7.3a with the following arguments: --outMultimapperOrder Random -- outSAMattrIHstart 0 --outFilterType BySJout --outFilterMismatchNmax 4 -- alignSJoverhangMin 8 --outSAMstrandField intronMotif --outFilterIntronMotifs RemoveNoncanonicalUnannotated --alignIntronMin 20 --alignIntronMax 1000000 -- alignMatesGapMax 1000000 --outWigType bedGraph --outWigNorm None -- outFilterScoreMinOverLread 0 --outFilterMatchNminOverLread 0 (Dobin et al., 2012). Reads per gene in the Gencodev33 gtf file were counted using the featureCounts function of Subread1.6.2 (Liao et al., 2013). This count matrix was used as input for DESeq2 to identify differentially expressed genes, calculating each time point versus the mock washout condition (Love et al., 2014). As reads mapping to the ERCC spike sequences were not different between conditions, the DESeq2 size factors were used to normalize samples.

### Western Blotting

Cells were rinsed with PBS, lysed in 2x Sample buffer (0.125 M Tris pH 6.8, 4% SDS, 20% Glycerol, 5% β-mercaptoethanol), sonicated and boiled at 95°C for 10 minutes. SDS-PAGE (Mini-Protean TGX, BioRad) in 0.025 M Tris, 0.2 M Glycine, 1% SDS (w/v) was followed by electrophoretic transfer to Protran nitrocellulose membrane (Cytiva) using the Mini Trans-blot wet-tank transfer system (BioRad) for 70 min at 250 mA in Transfer Buffer (0.02 M Tris, 0.223 M Glycine, 20% methanol). Membranes were stained with Ponceau Red (Fluka) to confirm successful transfer and blocked in 5% non-fat dry milk in TBS-Tween buffer (TBS-T; 20 mM Tris pH 7.6, 150 mM NaCl, 0.1% Tween-20) at room temperature. Incubations with primary and secondary antibodies were performed in TBS-T containing 5% non-fat dry milk. Signals were detected using an Odyssey CLx System (Li-Cor).

### Flow Cytometry

Cells were rinsed with PBS and detached from cultured plates using 0.5 mM EDTA in PBS for 5 min at 37°C and centrifuged for 4 min at 233 *g*. Cell pellets were resuspended by addition of ice-cold PBS supplemented with 2% FBS and counted using a TC-20 cell counter (BioRad). 2.5-5x10^5^ cells were harvested, spun down (400 *g*, 3 min, 4°C), and dissolved in 2% FBS in PBS containing 2.5 µl antibody. Antibody incubation was performed for 1 hour at 4°C in the dark. Labeled cells were then washed 3 times with 500 μl 2% FBS/PBS and centrifuged for 3 min at 400 *g* and 4°C. Cell pellets were dissolved in 2% FBS in PBS and flow cytometry was performed using an Accuri C6 Plus (BD Biosciences) or Citole Flow Cytometer (Beckman Coulter).

### Luciferase Notch reporter assay

0.8x10^5^ SVG-A receiver cells were seeded in each well of a 24-well plate. The following day, cells were transfected with 49 ng of a TP1-Luciferase (Kurooka et al., 1998; Minoguchi et al., 1997) and 1 ng Renilla-Luciferase (pRL-TK, Promega) using Lipofectamine™ 2000 (Invitrogen) according to manufacturers instructions. 24 hours after seeding, cells were either left untreated, treated with a γ- secretase inhibitor (GSI; Compound E at 0.5 μM), an ADAM10 inhibitor (GI254023X at 5 μM) or ligand blocking antibodies (see key resource table for concentrations). At this time, 1x10^5^ sender cells were added to each well after they were detached from a TC dish using 0.05% Trypsin/0.53 mM EDTA (Corning), and counted using a TC-20 cell counter (BioRad). Approximately 24 hours after co-culture, cells were rinsed with PBS, and lysed with 133 μl 1xPLB (Passive Lysis Buffer; Dual-Luciferase Reporter Assay System, Promega). 10 μl of each sample was analyzed using a GloMax Discover Microplate Reader (Promega) with 50 μl LARII (Luciferase Assay Reagent; Promega) and 25 μl Stop&Glo solution supplemented with the Stop&Glo substate (Promega).

### Calibration of HaloTag^JFX549^ in solution

The concentration of N2-C (*i.e.* NICD) in the nucleus of SVG-A (NOTCH2) cells was estimated by using a calibration method based on 3D imaging of recombinant HaloTag protein (rHaloTag) coupled to JFX549 in solution using spinning disk confocal microscopy. rHaloTag was expressed in *E. coli*, purified as described (Wilhelm et al., 2021) and labeled with JFX549 (the fluorophore used for visualization of N2-C). Specifically, 2 μM of rHaloTag was labeled with 8 μM JFX549 (50 u, ∼ 4x molar excess) in buffer solution (50 mM HEPES pH 7.3, 150 mM NaCl) for 25 min at room temperature in a total volume of 100 μl. A Zeba^TM^ Spin Desalting Column (7K MWCO; Thermo Scientific), pre-washed three times with 100 μl of buffer solution by centrifugation for 1 min at 1500 *g*, was used to remove unbound JFX549 ligand. Then, rHaloTag-JFX549 was applied to the column and centrifuged for 1 min at 1500 *g*. The flow-through was collected, the amount of rHaloTag determined by absorbance at 280 nm while the amount of JFX549 was determined by absorbance at 549 nm using a NanoDrop spectrophotometer (Thermo Scientific). A fluorescence calibration curve was then established by correlating the fluorescence intensity (F.I.) of solutions with different concentrations of rHaloTag^JFX549^ in imaging media using the spinning disk confocal microscope. Specifically, Z-stacks of 30 planes with 0.7 μm spacing between each optical plane and exposure time of 100 ms (561 nm laser) were acquired. Fluorescence intensity values from all planes were averaged and the background values obtained from imaging of the imaging media alone was subtracted. Calibration curves were obtained by fitting a linear equation to the experimental data acquired with the CCD (QuantEM, 512SC, Photometrics) or sCMOS (Prism 95B, Teledyne Photometrics) cameras (Figure S9A,B).

### HaloTag and DNA labeling

Cells were rinsed in imaging medium (Fluorobrite DMEM; Gibco) supplemented with 5% FBS (GeminiBio), 25 mM HEPES pH 7.4 (Gibco), 2 mM GlutaMax (Gibco), and 100 U/ml penicillin and streptomycin (Gibco). Cells were subsequently incubated at 37°C for 15 min with 100 nM JaneliaFluor dye (JFX549 or JFX646, gift from Luke Lavis, Janelia Research Campus) dissolved in imaging medium, then rinsed three times with imaging medium before bathing in fresh imaging medium used during imaging. Unlabeled knockin cells, unlabeled parental cells, or JFX-labeled parental cells were imaged as controls to evaluate the specificity of HaloTag labeling. Nuclear DNA was labeled by incubating the cells for 15 min at 37°C with SiR-DNA (1:4000; Spirochrome) in imaging media during or after HaloTag labeling.

### Cell delivery and cell pairing during imaging using spinning disk confocal microscopy

1.5x10^4^ SVG-A cells were seeded onto 8-well cover slips (Cellvis, C8-1.5H-N) to reach 30-50% confluency at the time of imaging. DMS53 cells were plated at a density of 6x10^4^ cells/well in a 24-well plate. Cells were incubated overnight at 37°C and 5% CO2 in DMEM supplemented with 10% FBS (GeminiBio, 100-106-500) and 100 U/ml penicillin and streptomycin (ThermoFisher Scientific, 15140163). The following day, plated SVG-A and DMS53 cells were labeled as described above with JFX549 and JFX646 dyes, respectively. For pairing, DMS53 cells were detached by incubation with PBS supplemented with 0.5 mM EDTA for 3 min at 37°C. Cells were transferred into 1.5 ml microcentrifuge tubes and the PBS/EDTA solution was removed by spinning down the cells for 5 min at 400-1000 *g*. The DMS53 cells were then resuspended in 200 μl imaging media and 150 μl of this solution was dispensed on top of SVG-A cells plated in the 8- well cover slips. 3D live spinning disk confocal imaging was then performed. Images of SVG-A cells, acquired before addition of the DMS53 sender cell suspension, were used as controls.

### Microfluidics device

The microfluidics devices were fabricated as previously described with some modifications (Salman et al., 2020; see Video S1). Briefly, photomasks were designed with AutoCAD (AutoDesk Corp.), printed by CAD/Art Services, Inc. and placed in a clean room on top of 76.2 mm silicon wafers (University Wafer, 447) to produce by photolithography 60 µm depth molds using SU-8 2050 photoresist (Microchem, now Kayaku Advanced Materials, Inc.). A 10:1 mixture of Sylgard 184 elastomer Polydimethylsiloxane (PDMS) and curing agent (Sylgard 184 silicone elastomer kit, Dow Corning) was freshly prepared, degassed for 30 min, then poured on top of the silicon wafer and spin-coated at 1000 rpm for 60 s to achieve 100 μm thickness. After degassing in vacuum for 10 min, the silicon wafer covered by the unpolymerized PDMS film was cured by incubation at 65°C for 24 hours, after which the PDMS film was peeled off and placed on top of lab tape inside a plastic petri dish. Above the sites at which the inlet / outlet tubing were later attached to the device, we placed a strip of 400-700 µm thick PDMS film bonded to the site using an air plasma cleaner (PDC-001 plasma cleaner, Harrick Plasma) at 700 mTorr, 30 W for 1.5 min followed by incubation at 60°C for 20 min. Afterwards, the PDMS film was flipped upside down and a 0.35 mm hole was punched at the tubing attachment sites using a Ted Pella puncher. The chips were plasma bonded to 25 mm diameter glass coverslips (CS-25R15 - 150 µm thickness, Glaswarenfabrik Karl Hecht) freshly cleaned by sonication for 15 min in 1M KOH followed by 3 washes in distilled water.

Tube connections to the chips were made by connecting and sealing (epoxy) polyurethane tubing of 0.007” ID x 0.14” OD (BTPU-014, Instech) into Tygon tubing of 0.010” ID x 0.030” OD (06419-00, Cole-Parmer). The polyurethane tubing was then connected to the microfluidic device and sealed with epoxy (Figure S5). Before use, the microfluidic devices were sterilized by first flowing 70% ethanol through the tubing and channels and then placing the device for 5 hours in 70% ethanol. Prior to cell plating, the ethanol was removed by 5 sequential rinses with sterile PBS.

### Spinning disk confocal microscopy

Cells were detached using trypsin, counted, and seeded onto 8-well cover slips (Cellvis, C8-1.5H-N) in imaging media at 37°C in presence of 5% CO2 at densities chosen to reach 30-50% confluency at the time of imaging the following day. Images were acquired using a Zeiss Axio-Observer Z1 (Zeiss) equipped with a 63x objective (Plan-Apochromat, NA 1.4, Zeiss), a spinning disk confocal head (CSU-XI, Yokogawa Electric Corporation) with additional system magnification of 1.2x, and a spherical aberration correction system (Infinity Photo-Optical) controlled with a Marianas system (3i, Intelligent Imaging Innovation). Volumetric images were collected with 0.7 μm spacing between each optical plane and fluorescence recorded with a CCD (QuantEM, 512SC, Photometrics, 0.212 x 0.212 μm/pixel in xy) or a sCMOS camera (Prim 95B, Teledyne Photometrics, 0.145 x 0.145 μm/pixel in xy). The fluorophores were excited using solid-state lasers (Coherent Inc.) with λ excitation at 488, 561, or 640 nm coupled to an acoustic-optical tunable filter or the LaserStack (3i, Intelligent Imaging Innovation) using solid state diode lasers coupled through single mode optical fibers to the LaserStream^TM^ (3i, Intelligent Imaging Innovation). With the CCD camera, exposure times of 100 ms in all channels were used to image membranes, Notch synapses, and nuclei; exposure times of 50 ms were used to image vesicles. With the sCMOS camera, exposure times of 60 ms were used to image signals in the 561 and 640 nm channels, exposure times of 100 ms were used in the 488 nm channel to image cell nuclei, and exposure times of 50 ms (488 nm channel), 30 ms (561 nm channel), and 60 ms (640 nm channel) were used to image vesicles.

### Lattice light-sheet microscopy modified with adaptive optics (MOSAIC)

Time-lapse live 3D z-stacks were acquired using a lattice light-sheet microscope modified with adaptive optics, referred here as MOSAIC (Multimodal Optical Scope with Adaptive Imaging Correction). Live cell volumetric imaging was achieved by acquiring single time points at 1 min intervals for 1 hour or longer. Sequential images spaced 0.40 μm between each plane along the z-imaging axis were obtained in sample scan mode; each time point consisted of z-stack comprised of 90-200 z-planes. Samples were illuminated with a dithered multi-Bessel lattice light-sheet (Chen et al., 2014) with 0.50 inner and 0.55 outer numerical apertures (NA) of the annular mask; lasers (MPB Communications Inc.) emitting at 488, 560 or 642 nm were used for illumination. A 0.65 NA (Special Optics) and a 1.0 NA objective (Zeiss) were used for illumination and detection using sCMOS cameras (Hamamatsu, ORCA Flash 4.0 v3) with 0.104 x 0.104 μm/pixel in xy for data visualization. Typical exposures were 50 ms for 488 nm (mNeonGreen – N2-N), 20 ms for 560 nm (HaloTag labeled with JFX549 – N2-C), and 20 ms for 642 nm (SiR-DNA or HaloTag labelled with JFX646 – DLL4).

### Cell delivery and cell pairing during imaging using MOSAIC

1.5x10^5^ SVG-A cells were plated onto the center of the microfluidics device, followed by overnight incubation at 37°C and 5% CO2 in DMEM supplemented with 10% fetal bovine serum (FBS, GeminiBio, 100- 106-500) and 100 U/ml penicillin and streptomycin (ThermoFisher Scientific, 15140163). Prior to imaging, the cells were labeled as described. The microfluidics device with attached SVG-A cells was then placed on the MOSAIC sample holder and its inlet tubing (Tygon tubing 0.010” ID x 0.030” OD (06419-00, Cole-Parmer)) was connected to the flow meter (Flow Unit M Flow-Rate Platform, Fluigent) (Figure S5). Another tubing, connected to a 2 ml microcentrifuge tube (Eppendorf) with an air-tight metal tube cap (P-CAP 2 mL High Pressure, Fluigent) containing a suspension of 5x10^5^ DMS53 cells labeled with JFX646, was also connected to the inlet of the flow meter. The sealed tube was further connected to the pressure controller (Microfluidic Flow Control System – EZ, Fluigent) using pneumatic tubing. The tube with suspended DMS53 cells was kept up to 5 min at 37°C (dry bath, My Block, Benchmark) before cell injection into the microfluidics device. Inlet pressure of 50-100 mbar and a flow of 10-15 μl/min for 30-90 s of the suspension containing DMS53 sender cells were controlled in real time using MAESFLO software (Fluigent). Upon ending the flow, the DMS53 cells were allowed to settle by gravity and to pair with the SVG-A cells attached in the microfluidics device.

### Fluorescence recovery after photobleaching (FRAP)

Fluorescence recovery after photobleaching (FRAP) was performed with the spinning disk confocal microscope by photobleaching a region of interest (ROI) of 1 μm in radius for 5 ms using 100% laser power. A 100 ms exposure time was used to collect images every 1 s for 10 s before bleaching and for 60 s after bleaching. SVG-A and DMS53 cells, alone or in pairs were used to perform single FRAP experiments for a given isolated cell or cell-pair. For photobleaching of synapses, 1-2 ROI were selected on the Notch synapse, while another ROI elsewhere on the cell membrane was used as a control. The position of the synapse within the ROI was determined by imaging in a non-bleached channel. A similar time series acquired in a different region of the cell not subjected to FRAP was used to correct for bleaching due to imaging only.

## Data analysis

### Ratiometric analysis

The relative amounts of N2-N (mNeonGreen), N2-C (HaloTag) and DLL4 (HaloTag) or JAG1 (HaloTag) associated with the Notch synapse, excluded from it and in the cell membrane, or associated with vesicles in the sender cell were determined by ratiometric analysis of the corresponding fluorescence signals within appropriate ROIs. The first step in the ratiometric analysis consisted in determining the relative amount of N2-N, N2-C and DLL4 (or JAG1) within a given image. This step was achieved by comparing the fluorescence intensity of N2-N with respect to N2-C (HaloTag^JFX549^) or N2- N with respect to DLL4 (HaloTag^JFX549^) or JAG1 (HaloTag^JFX549^). The second step established the relative signal resulting from JFX549 and JFX646 labeling by comparing the relative fluorescence intensity of N2-C (HaloTag^JFX549^) in one sample with respect to N2-C (HaloTag^JFX646^) in a second independently labeled sample.

Ratiometric analysis of fluorescence signals within appropriate ROIs was performed by using a Macro written for Fiji (Schindelin et al., 2012). A three-pixel width line was drawn across the membrane or synapse as an ROI to obtain the mean fluorescence intensity (F.I.) of the measured values within the line width. To account for the three dimensionality, two planes below and two planes above the ROI were determined, resulting in 5 ROI, one per plane. The maximum intensity F.I. in those 5 ROI (usually coincident with the main ROI) was subtracted by the F.I. of the background to obtain the F.I.^max^, a value proportional to the density of molecules at the Notch synapse, on the surrounding cell membrane surface, and in vesicles within the sender cell.

The ratiometric quantification of N2-N associated with intracellular vesicles in the DMS53 sender cell was corrected by the intrinsic autofluorescence F.I.^af^, determined in unpaired DMS53 cells imaged in the microfluidics device 60 minutes after initiation of the cell pairing experiment.

### FRAP analysis

FRAP analysis was conducted as described (Govindaraj & Post, 2021) using Fiji (Schindelin et al., 2012) on the fluorescent signal within the photobleached ROI of the Notch synapse or plasma membrane after correcting the fluorescent signals for the inherent photobleaching due to imaging; the fluorescence intensity of the first 10 time points prior to FRAP were averaged and normalized to 1. The FRAP recovery curve was fitted using a single decay exponential from which the diffusion coefficient was estimated as D = (0.224 x r^2^)/ t1/2, where r is the radius of the bleached ROI and t1/2 the half-life of recovery (Kang et al., 2012).

### Nuclear N2-C (i.e. NICD) concentration

The nuclear N2-C (*i.e.* NICD) concentration was estimated by applying the volume calibration curve to the mean nuclear NICD fluorescence intensity (F.I.) from the non-punctate and diffuse nuclear N2-C signal. A macro written using Fiji (Schindelin et al., 2012) was used to automate the calculations. A binary mask of the nucleus defined by the SiR-DNA signal from Notch cells was used to define the nuclear region from which to calculate the averaged intensity per plane; an estimate of the nuclear volume was obtained by multiplying the z-planes by the space between optical planes (0.7 μm). The extent of out of focus fluorescence contributed by molecules located on the plasma membrane to different z-planes within the nucleus was estimated by measuring the fluorescence of N2-C (*i.e.* NECD; which is always absent from the nucleus). This value was then used to correct for the contribution of out of plane N2-C signal from the plasma membrane to the nuclear signal value.

### Nuclear N2-C (i.e. NICD) concentration, N2-N molecules in synapse and in DMS53 cell vesicles in the time-lapse 3D z-stacks acquired using MOSAIC

The fluorescence signals obtained with MOSAIC were normalized to the signals obtained with the SD. This normalization was done by determining the ratio of N2-C (HaloTag^JFX549^) fluorescence within a plane orthogonal to the plasma membrane acquired with MOSAIC and SD. The nuclear N2-C concentration was estimated as above using SD.

A binary mask corresponding to the Notch synapse was defined by the logical intersection of the N2-N, N2-C and DLL4 signals. The averaged N2-N fluorescence signal per pixel (0.1x0.1x0.4 μm) times the number of pixels corrected by N2-N membrane signal outside of the synapse and normalized by the signal ratio between N2-N and N2-C on the membrane corresponds to the number of N2-N (*i.e.* NECD) molecules in the synapse. Vesicles containing N2-N in DLL4 cells were identified using the 3D cmeAnalysis software (Aguet et al., 2016). The volume of a given vesicle was defined as a box of 3x3x3 (x,y,z) pixels from which the N2-N average fluorescence and the number of molecules per vesicle were calculated as described above. This number multiplied by the number of vesicles corresponded to the total amount of N2-N transendocytosis into the DLL4 cell.

### Statistical analysis

All statistical analyses were performed using GraphPad Prism (GraphPad). Sample distribution and normality tests were performed for each data set. Statistical tests that were used are indicated in the figure legends.

### Data and code availability

Raw data, MATLAB codes, and FIJI macros are available upon request. RNA-Seq data is accessible at NCBI GEO database (Edgar et al., 2002) accession GSE235637.

## Figure legends

**Figure S1.**
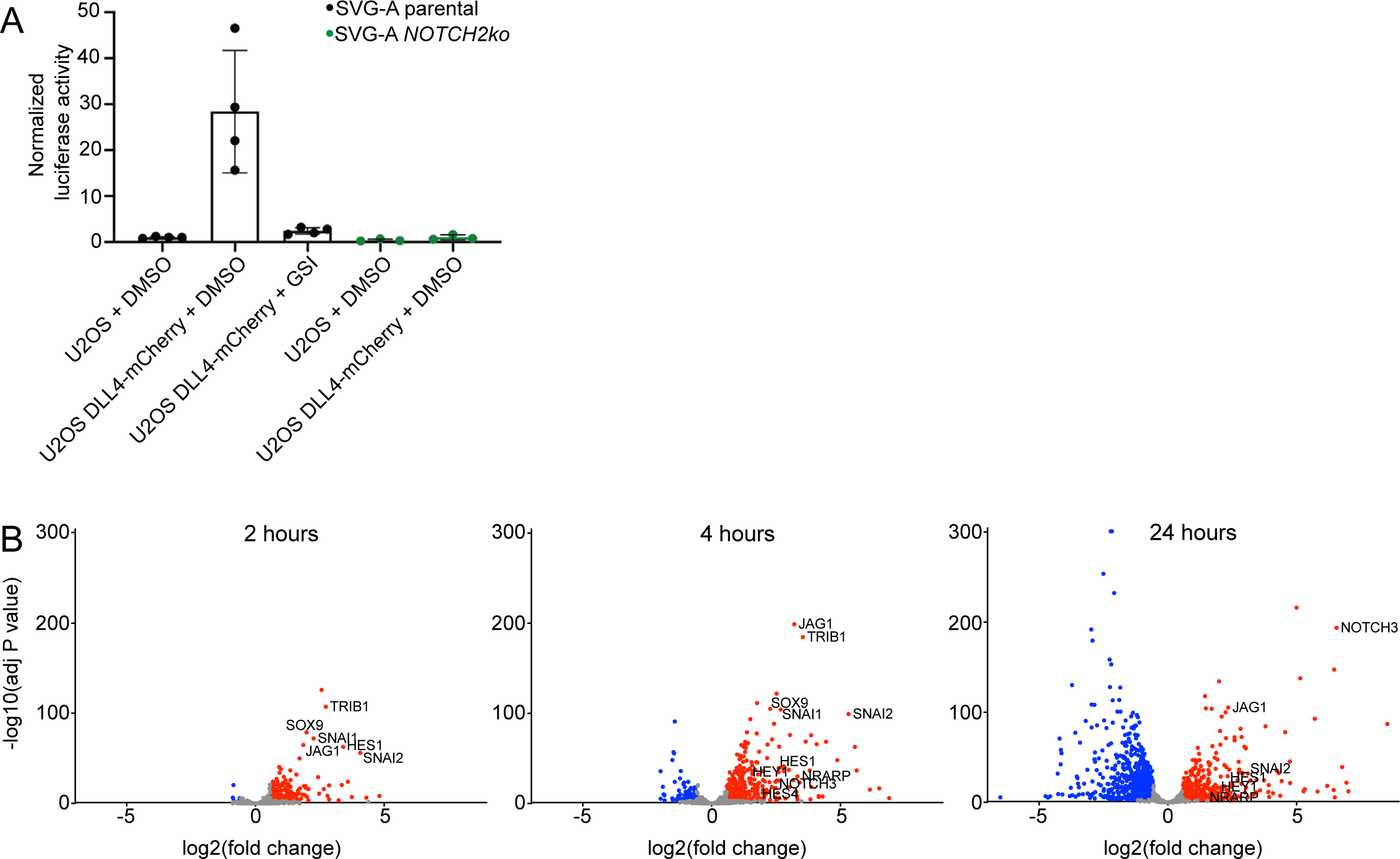
SVG-A cells as Notch receiving cells. **A.** Luciferase reporter gene assay for Notch-induced transcription (Malecki et al., 2006). Parental and *NOTCH2ko* SVG-A cells were co-transfected with a reporter plasmid containing firefly luciferase under control of the Notch-responsive TP1 promoter (Kurooka et al., 1998; Minoguchi et al., 1997) and an internal control Renilla luciferase plasmid (Promega) using Lipofectamine™ 2000 (Invitrogen). 24 hours after transfection, these cells were co-cultured with U2OS parental cells or U2OS cells stably expressing DLL4-mCherry in presence of DMSO or GSI (Compound E). Cells were lysed 24 hours later, and the firefly and Renilla luciferase activity of each lysate was measured using a Dual Luciferase Assay Kit (Promega). The firefly/Renilla ratio was normalized to the signal for co-culture of SVG-A cells with parental U2OS cells. Plots show mean ± standard deviation from four independent biological replicates (n=4). **B.** RNA-seq analysis of genes induced in SVG-A cells seeded onto tissue culture plates coated with JAG1-Fc (200 μg/ml) at timepoints after removal of GSI (100 nM). Volcano plots compare RNA abundance at 2, 4, and 24 hours to a reference sample at t=0 subjected to a mock washout with media containing GSI. Red dots indicate significantly upregulated genes (adj. p value < 0.001, Fold Change > 1.5), while blue dots indicate significantly downregulated genes (adj. p value < 0.001, Fold Change < -1.5).

**Figure S2.**
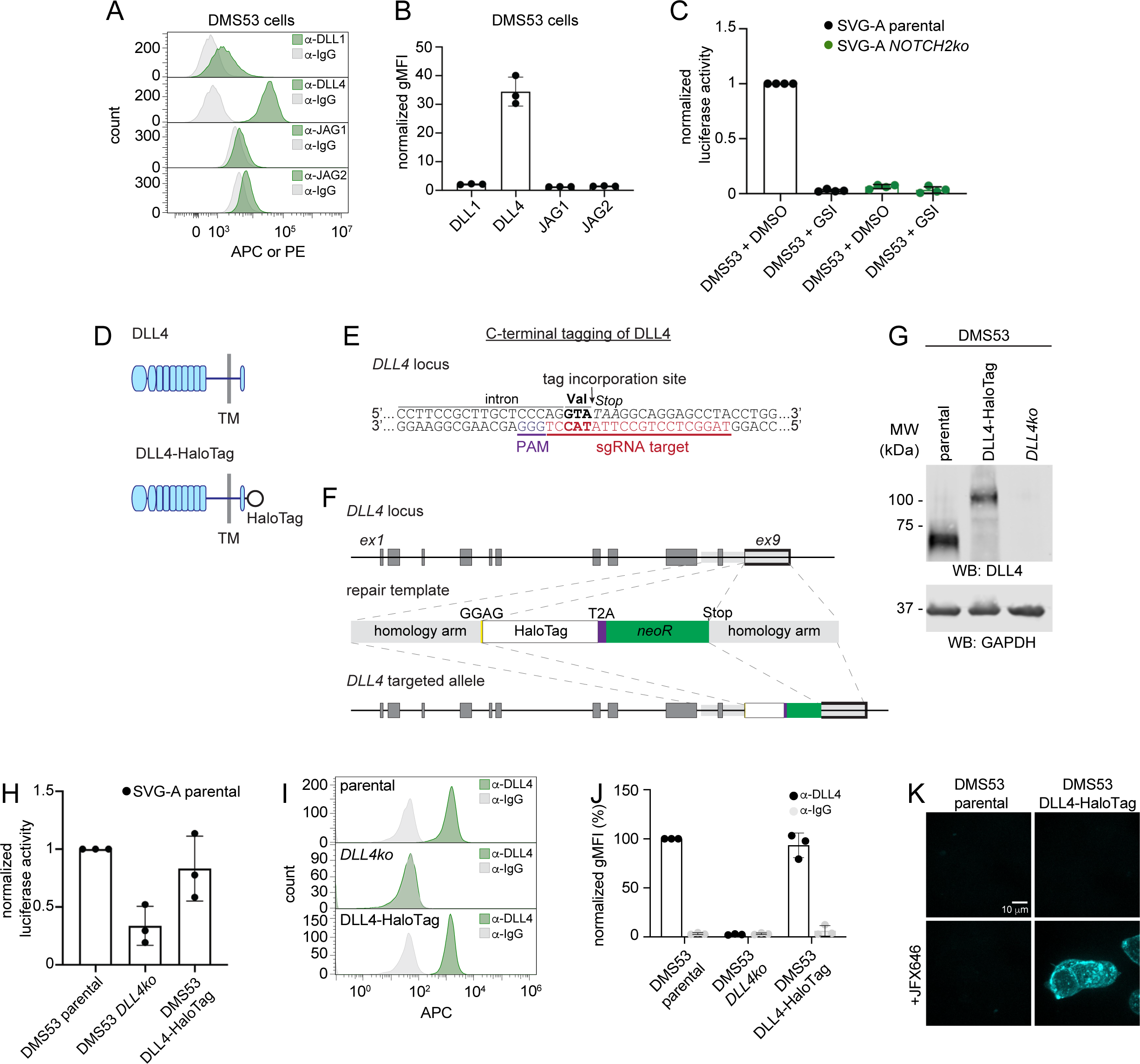
Preparation of DMS53 DLL4-HaloTag knockin cells. **A.** Analysis of Notch ligands on DMS53 cells by flow cytometry. Cells were stained using fluorescently conjugated anti-DLL1, anti-DLL4, anti-JAG1, and anti-JAG2 antibodies, using anti-IgG isotype staining as a reference. Representative histograms of stained cell populations are shown. **B.** Geometric mean fluorescence intensities (gMFI) of anti-DLL1, anti-DLL4, anti-JAG1, and anti-JAG2 stained cell populations analyzed by flow cytometry in **A**, normalized to anti-IgG isotype staining. **C.** Luciferase reporter gene assay. Parental or *NOTCH2ko* SVG-A cells were co-transfected with a reporter plasmid containing firefly luciferase under control of the Notch-responsive TP1 promoter (Kurooka et al., 1998; Minoguchi et al., 1997) and an internal control Renilla luciferase plasmid (Promega) using Lipofectamine™ 2000 (Invitrogen). 24 hours after transfection, these cells were co-cultured with DMS53 cells in the presence of DMSO or GSI (0.5 μM). Cells were lysed 24 hours later, and the firefly and Renilla luciferase activity of each lysate was measured using a Dual Luciferase Assay Kit (Promega). The firefly/Renilla ratio was normalized to the signal for co-culture of SVG-A parental cells with DMS53 cells in the presence of DMSO. **D.** Schematic showing parental DLL4 and the engineered DLL4 fusion containing a C-terminal HaloTag. **E.** Sequence from the *DLL4* locus showing the C-terminal tagging site and sgRNA targeting sequence used for CRISPR/Cas9 mediated genome editing. **F.** *DLL4* genomic locus showing schematics of the repair template with homology arms, tagging site and linker used, as well as the targeted *DLL4* allele. **G.** Anti-DLL4 western blot. Lysed parental DMS53 cells, *DLL4-HaloTag* knockin DMS53 cells, and DMS53 *DLL4*ko cells were probed with an anti-DLL4 antibody. GAPDH immunodetection was used as a loading control. **H.** Luciferase reporter gene assay. SVG-A cells were treated as in **C** before co-culturing with DMS53 parental, DMS53 *DLL4ko*, or DMS53 DLL4- HaloTag cells. Cells were lysed 24 hours later, and the firefly and Renilla luciferase activity of each lysate was measured using a Dual Luciferase Assay Kit (Promega). The firefly/Renilla ratio was normalized to the signal for co-culture of SVG-A cells with parental DMS53 cells. **I, J.** Flow cytometry analysis of DMS53 parental, DMS53 *DLL4ko*, and DMS53 DLL4-HaloTag cell lines. Cells were incubated with a fluorescently conjugated anti-DLL4 antibody or an anti-IgG isotype control. Representative histograms (**I**) and geometric mean fluorescence intensity plots, normalized to parental cells (**J**), are shown. **K.** Imaging of DMS53 parental cells and DMS53 DLL4-HaloTag knockin cells. Cells were imaged using a spinning disk confocal microscope when unlabeled or when labeled with JaneliaFluorX646 (JFX646). Scale bar as indicated. Data plotted in **B, C, H, and J** are shown as mean ± standard deviation, with n ≥ 3 independent biological replicates.

**Figure S3.**
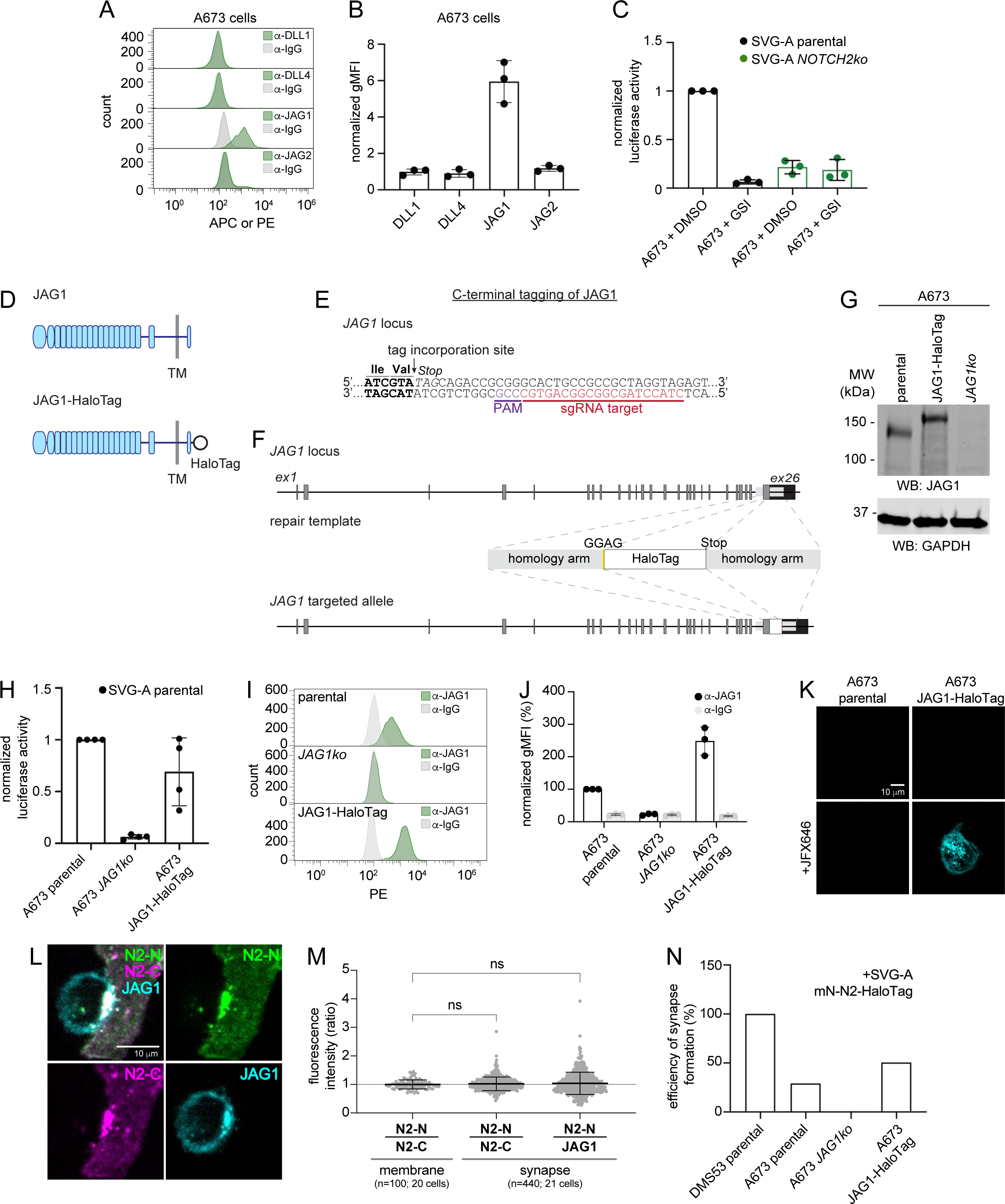
Preparation of A673 JAG1-HaloTag knockin cells. Analysis of Notch ligands on A673 cells by flow cytometry. Cells were stained using fluorescently conjugated anti-DLL1, anti-DLL4, anti-JAG1, and anti-JAG2 antibodies, using anti-IgG isotype staining as a control. Representative histograms of staining are shown. **B.** Geometric mean fluorescence intensities (gMFI) of anti-DLL1, anti-DLL4, anti-JAG1, and anti-JAG2 stained cell populations analyzed by flow cytometry in **A**, normalized to anti-IgG isotype staining. **C.** Luciferase reporter gene assay. Parental or *NOTCH2ko* SVG-A cells were co-transfected with a reporter plasmid containing firefly luciferase under control of the Notch-responsive TP1 promoter (Kurooka et al., 1998; Minoguchi et al., 1997) and an internal control Renilla luciferase plasmid (Promega) using Lipofectamine™ 2000 (Invitrogen). 24 hours after transfection, these cells were co-cultured with A673 cells in the presence of DMSO or GSI (0.5 μM). Cells were lysed 24 hours later, and the firefly and Renilla luciferase activity of each lysate was measured using a Dual Luciferase Assay Kit (Promega). The firefly/Renilla ratio was normalized to the signal for co-culture of SVG- A parental cells with A673 cells in the presence of DMSO. **D.** Schematic showing parental JAG1 and the engineered JAG1 fusion containing a C-terminal HaloTag. **E.** Sequence from the *JAG1* locus showing the C-terminal tagging site and sgRNA targeting sequence used for CRISPR/Cas9 mediated genome editing. **F.** *JAG1* genomic locus showing schematics of the repair template with homology arms, tagging site and linker used, as well as the targeted *JAG1* allele. **G.** Anti-JAG1 western blot. Lysed parental A673 cells, JAG1-HaloTag knockin A673 cells, and A673 *JAG1ko* cells were probed with an anti-JAG1 antibody. GAPDH immunodetection was used as a loading control. **H.** Luciferase reporter gene assay. SVG-A cells were treated as in **C** before co-culturing with A673 parental, A673 *JAG1ko*, or A673 JAG1-HaloTag cells. Cells were lysed 24 hours later, and the firefly and Renilla luciferase activity of each lysate was measured using a Dual Luciferase Assay Kit (Promega). The firefly/Renilla ratio was normalized to the signal for co-culture of SVG-A cells with parental JAG1 cells. **I, J.** Flow cytometry analysis of A673 parental, A673 *JAG1ko*, and A673 JAG1-HaloTag cell lines. Cells were incubated with a fluorescently conjugated anti-JAG1 antibody or an anti-IgG isotype control. Representative histograms (**I**) and geometric mean fluorescence intensity plots, normalized to parental cells (**J**), are shown. **K.** Imaging of A673 parental cells and A673 JAG1-HaloTag knockin cells. Cells were imaged using a spinning disk confocal microscope when unlabeled or when labeled with JaneliaFluorX646 (JFX646). Scale bar as indicated. Data plotted in **B, C, H, and J** are shown as mean ± standard deviation, with n ≥ 3 independent biological replicates. **L.** Representative images of a synapse formed by pairing an mNeon-NOTCH2-HaloTag (labeled with JFX549) SVG-A knockin cell with a JAG1-HaloTag (labeled with JFX646) knockin A673 cell. N2-N is represented in green, N2-C in magenta, and JAG1 in cyan. Scale bars as indicated. **M.** Ratios of fluorescence intensities of signals associated with N2-N and N2-C in the membrane and of N2-N and N2-C or N2-N and JAG1 in synapses, respectively. Data plotted are shown as mean ± standard deviation. The number of synapses (n) and the number of cells analyzed are indicated. Statistical analysis was performed using Kruskal-Wallis one-way ANOVA; ns = p>0.05. **N.** Efficiency of synapse formation. SVG-A mNeon-NOTCH2-HaloTag cells were paired with DMS53 (DLL4) or different engineered forms of A673 (JAG1) cells.

**Figure S4.**
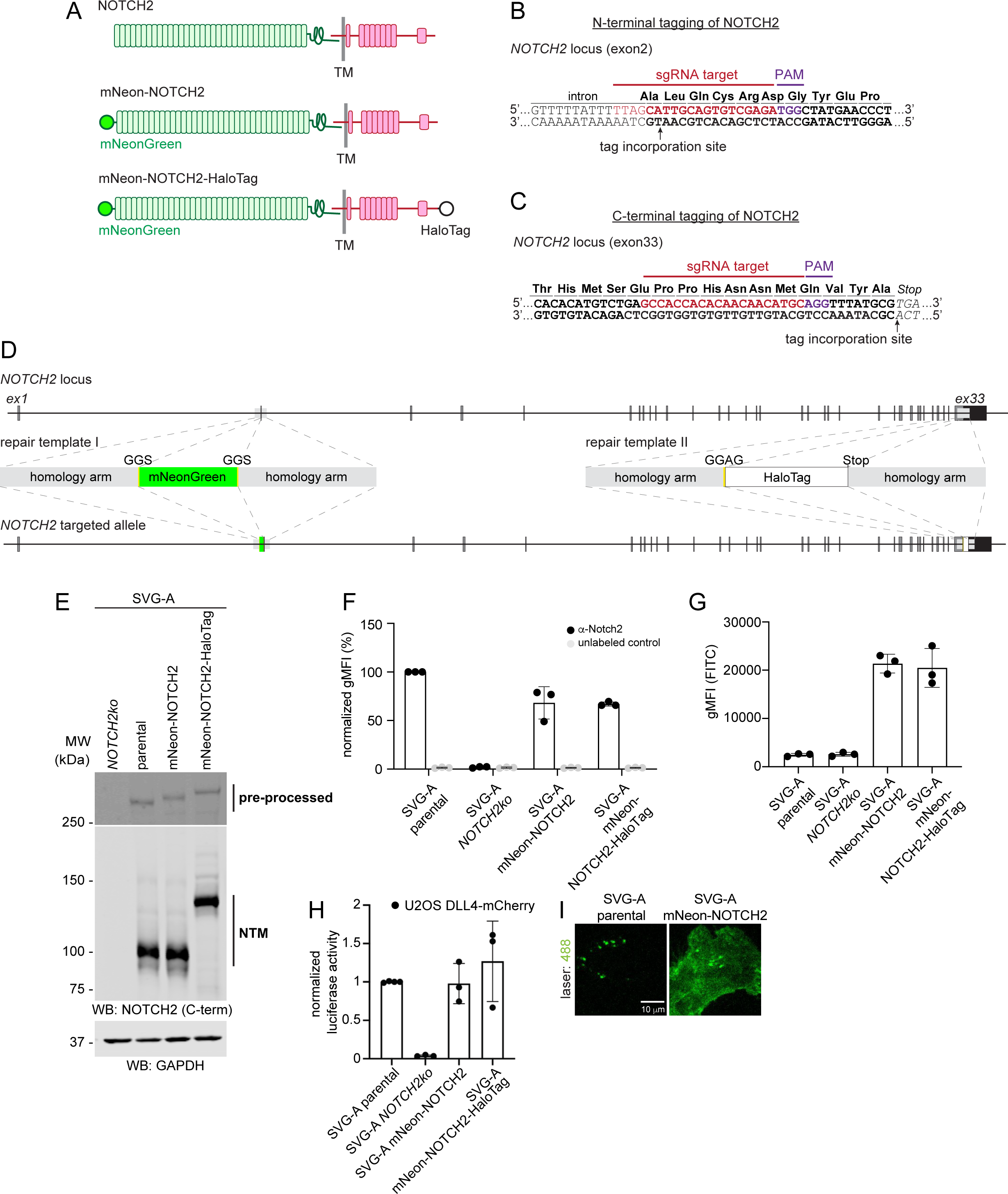
Tagging and screening of SVG-A NOTCH2 knockin cells. **A.** NOTCH2 domain organization and tagging sites. Untagged NOTCH2 (top), NOTCH2 N-terminally tagged with mNeonGreen (mNeon-NOTCH2) (middle), and NOTCH2 tagged with mNeonGreen at the N-terminus and a HaloTag at the C-terminus (mNeon-NOTCH2- HaloTag) (bottom) are shown. **B-D.** CRISPR/Cas9 genome editing at the N- and C-termini of NOTCH2. **B.** N-terminal tagging site and sgRNA targeting sequence. **C.** C-terminal tagging site and sgRNA targeting sequence. **D.** Schematic of *NOTCH2* locus, the repair templates with homology arms, tags, and linkers used, as well as the *NOTCH2* targeted allele. **E.** Western Blot of SVG-A cells with *NOTCH2* knockout (*NOTCH2ko*), parental cells, cell clone with N-terminal knockin of mNeonGreen (mNeon-NOTCH2), and the doubly tagged cell clone expressing NOTCH2 with an N-terminal mNeonGreen tag and a C-terminal HaloTag (mNeon-NOTCH2-HaloTag). The antibody recognizing the intracellular domain of NOTCH2 was used to detect the C-terminal NTM fragments of the heterodimeric receptors (middle). Detection of pre-processed proteins with the same antibody (top, longer exposure) confirms the integration of both tags. GAPDH immunodetection (bottom) was used as a loading control. **F, G.** Flow cytometry analysis of parental, *NOTCH2ko* and tagged clones. In **F**, cells were stained with an anti-NOTCH2- APC antibody (black) or unlabeled (gray), and in **G**, the mNeonGreen signal was detected in the FITC channel. **H.** Luciferase reporter gene assay. Parental, *NOTCH2ko*, and NOTCH2 knockin SVG-A cells (as indicated) were co-transfected with a reporter plasmid containing firefly luciferase under control of the Notch-responsive TP1 promoter (Kurooka et al., 1998; Minoguchi et al., 1997) and an internal control Renilla luciferase plasmid (Promega) using Lipofectamine™ 2000 (Invitrogen). 24 hours after transfection, these cells were co-cultured with U2OS cells stably expressing DLL4-mCherry. Cells were lysed 24 hours later, and the firefly and Renilla luciferase activity of each lysate was measured using a Dual Luciferase Assay Kit (Promega). The firefly/Renilla ratio was normalized to the signal for co-culture of SVG-A parental cells with U2OS DLL4-mCherry cells. **I.** Fluorescence of mNeonGreen-NOTCH2 cells (right) compared to parental cell autofluorescence (left), imaged on a spinning disk confocal microscope. Scale bar as indicated. Plots in **F-H** show mean ± standard deviation from at least three independent biological replicates (n≥3).

**Figure S5.**
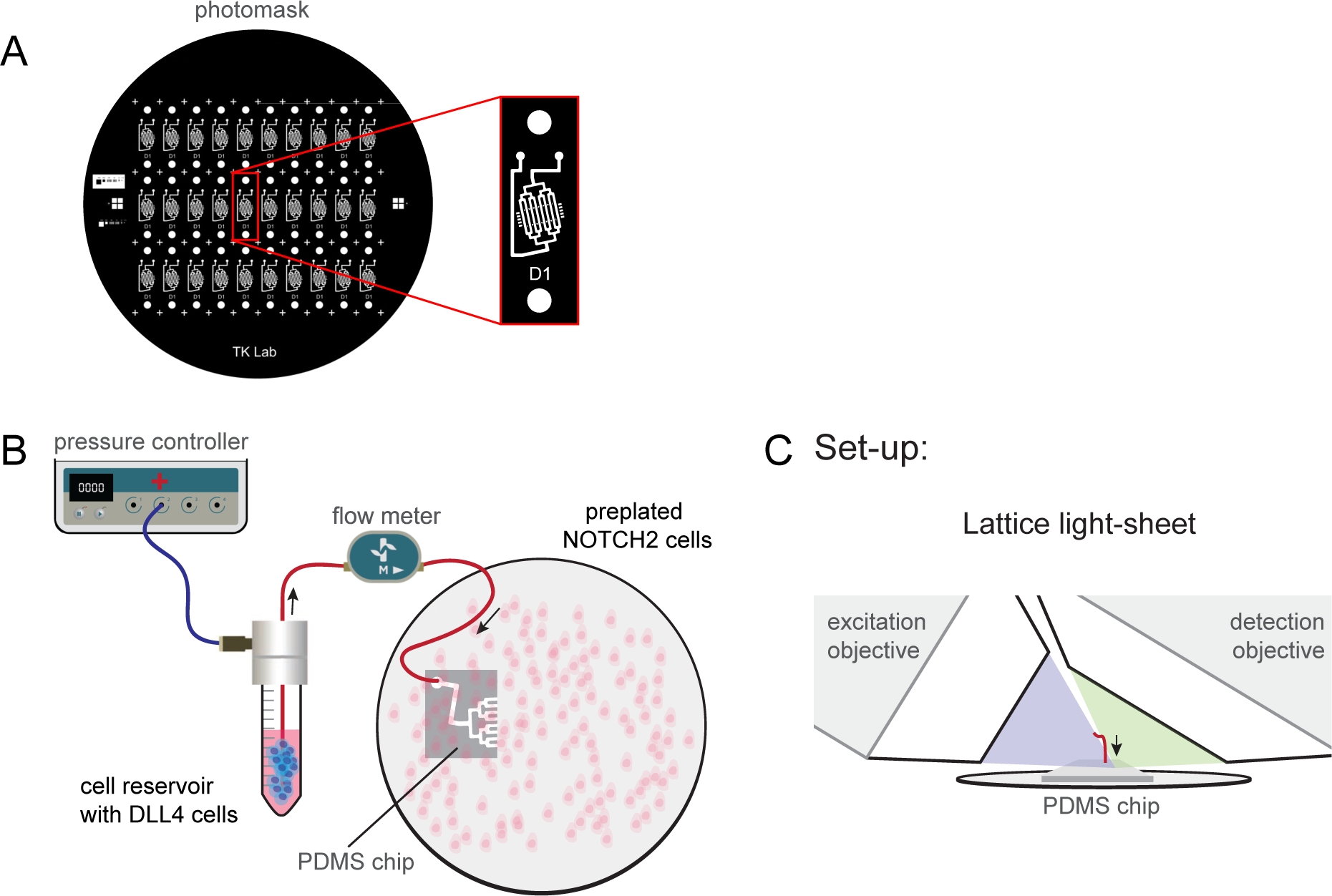
Microfluidics configuration for cell pairing. **A.** Scheme of the photomask as used to create the PDMS chips for microfluidic cell delivery. **B.** Schematic of the microfluidics system used to initiate pairing of DLL4 (DMS53) cells with NOTCH2 (SVG- A) cells. NOTCH2 cells were seeded on a coverslip containing a PDMS chip that is connected to a cell reservoir, and labeled with a JFX dye. DLL4 cells were labeled separately with a different JFX dye, detached from a culture dish, and stored in the cell reservoir until used for pairing. Using a pressure-based controller, the DLL4 cells were delivered to the pre-plated NOTCH2 cells. A flow meter was always used to monitor the flow rate. Images of the pressure controller, tube cap, and flow meter were adapted from the Fluigent Image package (Fluigent Microfluidics^TM^, Le Kremlin-Bicêtre, France). **B.** Positioning of the coverslip and PDMS chip when used for lattice light-sheet microscopy.

**Figure S6.**
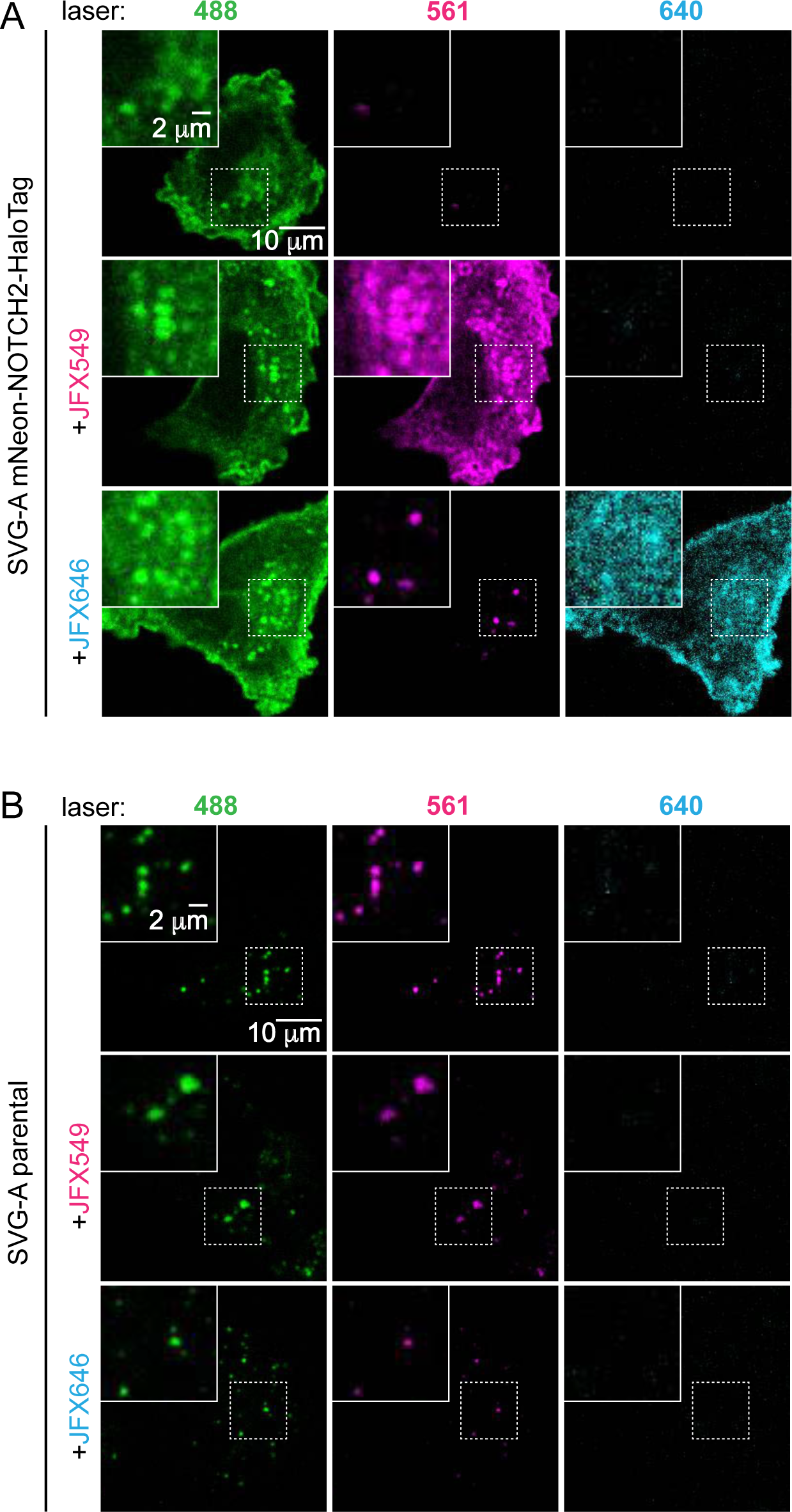
Fluorescence of unlabeled and dye-coupled SVG-A mNeon-NOTCH2- HaloTag cells. **A, B.** Representative images of SVG-A mNeon-NOTCH2-HaloTag cells (**A**) or parental SVG-A cells (**B**) when unlabeled, incubated with JFX549 or incubated with JFX646, acquired using 488, 561, and 640 lasers in a spinning disk confocal microscope. Images show the maximum intensity projection of z-stacks. Insets in the left upper corner are magnifications of the dashed square regions. Scale bars as indicated.

**Figure S7.**
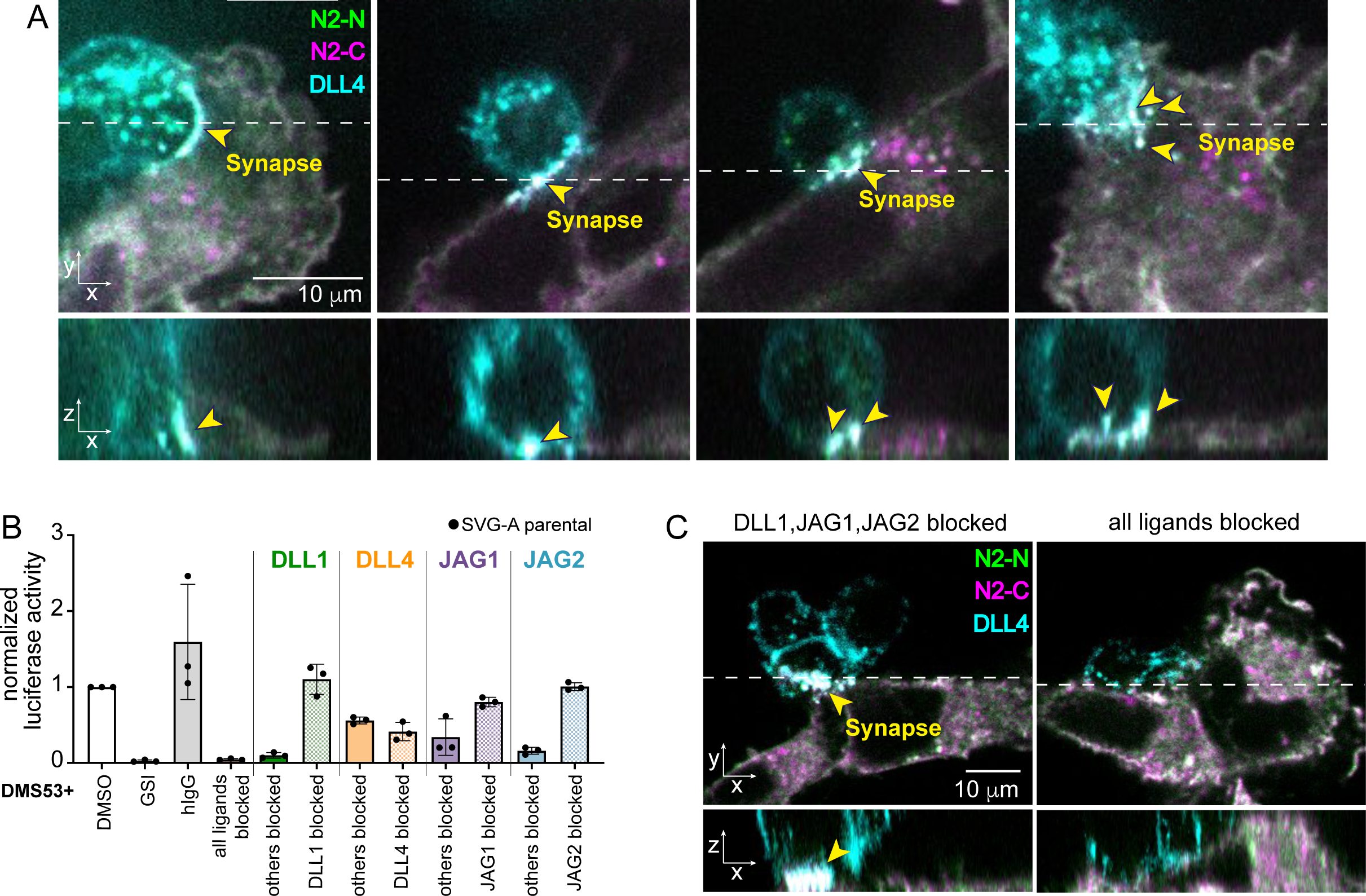
Synapse morphology and ligand dependence. **A.** Set of images representing the variability of synapses that form at sites of contact between DLL4 and NOTCH2 cells. Images show single planes of top and orthogonal views (dashed lines indicate the region used for the orthogonal views) acquired using a spinning disk confocal microscope. N2-N is in green, N2-C in magenta, and DLL4 in cyan. **B.** Luciferase reporter gene assay. Parental SVG-A cells were co-transfected with a reporter plasmid containing firefly luciferase under control of the Notch-responsive TP1 promoter (Kurooka et al., 1998; Minoguchi et al., 1997) and an internal control Renilla luciferase plasmid (Promega) using Lipofectamine™ 2000 (Invitrogen). 24 hours after transfection, these cells were co-cultured with DMS53 cells in the presence of DMSO or GSI (0.5 μM), human IgG (hIgG) antibody control, or different combinations of ligand-blocking antibodies. Cells were lysed 24 hours later, and the firefly and Renilla luciferase activity of each lysate was measured using a Dual Luciferase Assay Kit (Promega). The firefly/Renilla ratio was normalized to the signal for co-culture of SVG-A parental cells with DMS53 cells in the presence of DMSO. **C.** Representative single plane spinning disk confocal images and orthogonal views (dashed lines indicate the region used for the orthogonal views) showing synapse formation between DLL4 and NOTCH2 cells when DLL1, JAG2, and JAG2 ligand-blocking antibodies are present. No synapses are formed when all ligands are blocked by antibodies. N2-N is in green, N2-C in magenta, and DLL4 in cyan. Data in **B** are plotted as mean ± standard deviation, n = 3 independent biological replicates. Synapses in **A** and **C** are indicated by yellow arrowheads; scale bars as indicated.

**Figure S8.**
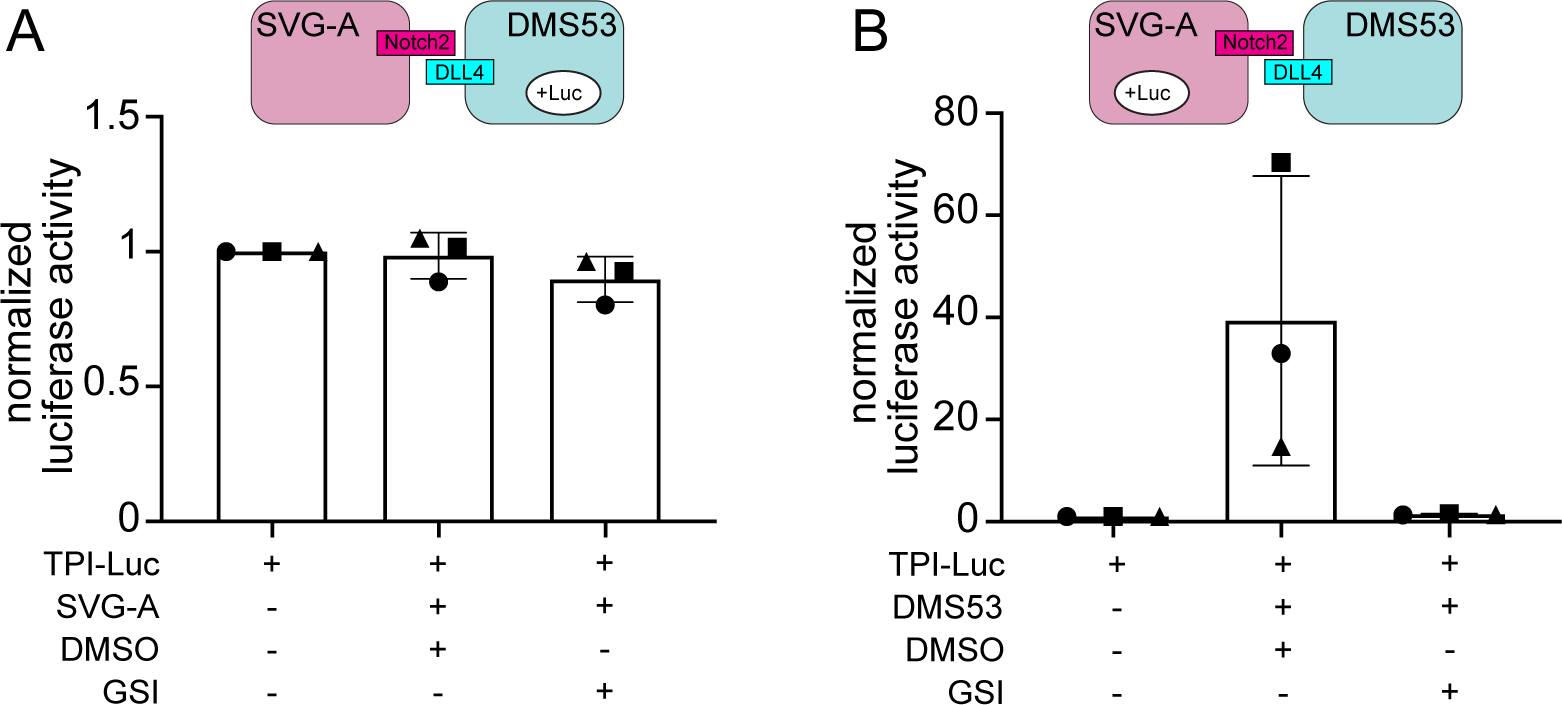
NOTCH2 signaling activity is not detected in DMS53 cells. **A, B**. Luciferase reporter gene assays. DLL4 (DMS53) (**A**) or NOTCH2 (SVG-A) cells (**B**) were co-transfected with a reporter plasmid containing firefly luciferase under control of the Notch-responsive TP1 promoter (Kurooka et al., 1998; Minoguchi et al., 1997) and an internal control Renilla luciferase plasmid (Promega) using Lipofectamine™ 2000 (Invitrogen). In **A**, Cells were cultured alone, co-cultured with NOTCH2 cells in DMSO, or co-cultured with NOTCH2 cells in presence of GSI (0.5 μM). In **B**, Cells were cultured alone, co-cultured with DLL4 cells in DMSO, or co-cultured with DLL4 cells in presence of GSI (0.5 μM). Cells were lysed 24 hours later, and the firefly and Renilla luciferase activity of each lysate was measured using a Dual Luciferase Assay Kit (Promega). The firefly/Renilla ratio was normalized to the signal for parental cells that did not undergo co-culture (left). Data are plotted as mean ± standard deviation, n = 3 independent biological replicates.

**Figure S9.**
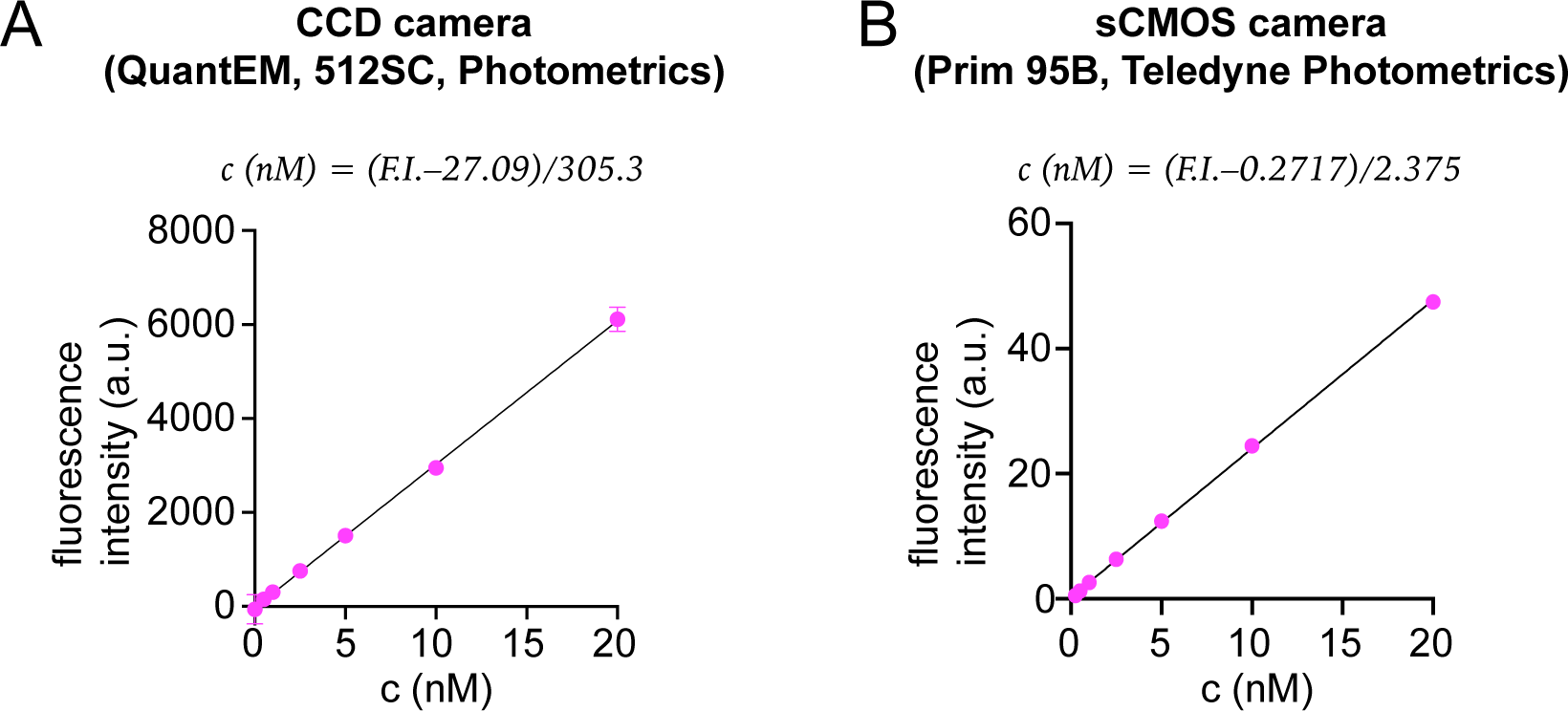
Calibration curves for determining the concentration of HaloTag labeled with JFX549 ligand. **A, B.** Plots of measured fluorescence intensity as a function of HaloTag-JFX549 concentration over a concentration range of 0-20 nM. A 100 ms exposure time was used in a CCD (**A**) or sCMOS (**B**) camera. Data are plotted as mean ± standard deviation. The equations representing the best fit line to the data are shown above each plot.

**Figure S10.**
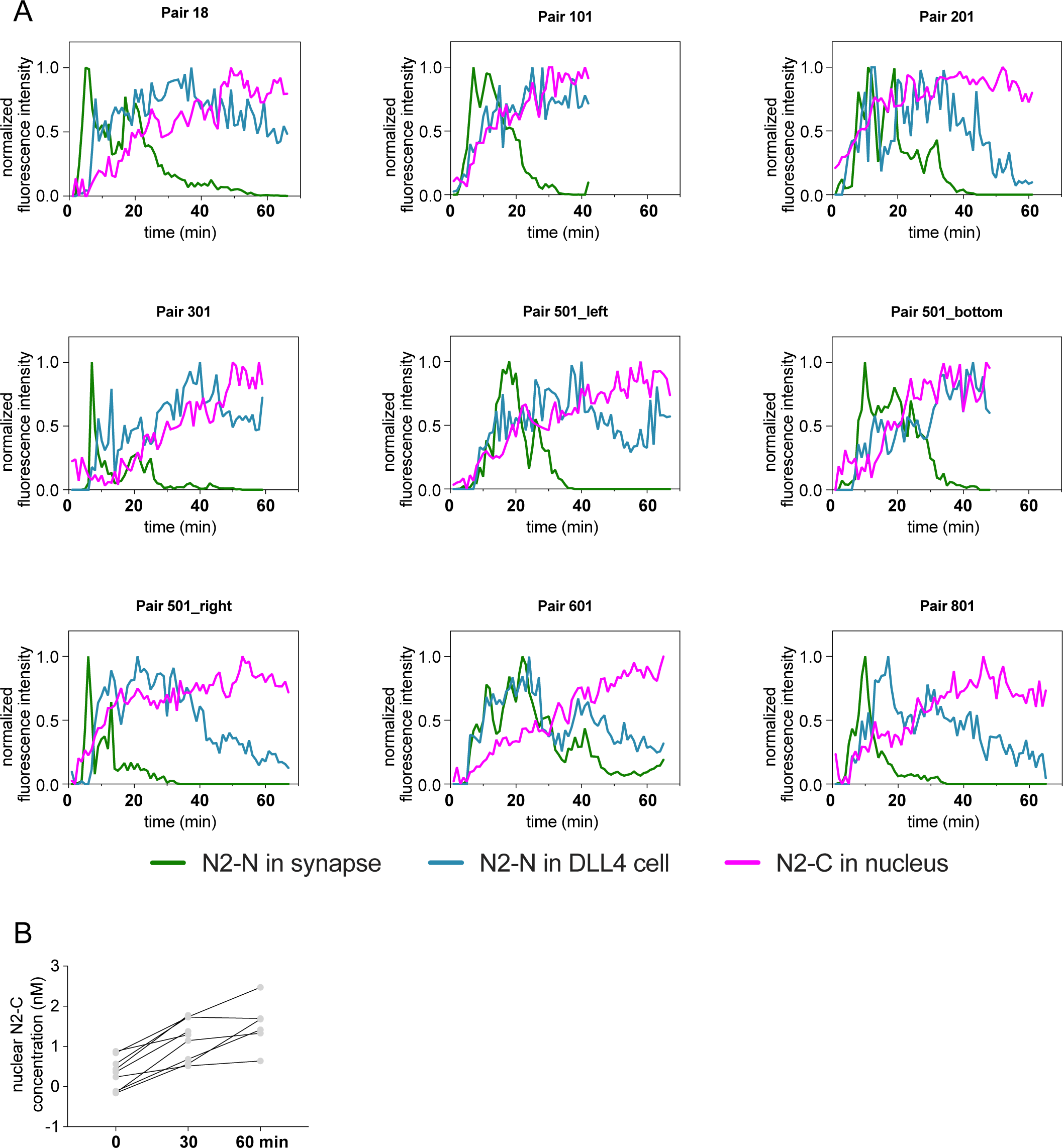
Plots of fluorescence as a function of time for nine independent cell pairing events. **A.** Plots showing the normalized fluorescence intensity of N2-N in synapses (green), N2-N in DLL4 cell vesicles (blue) and N2-C in nuclei of NOTCH2 cells (magenta) as a function of time after DLL4 cell contact. Graphs show n = 9 independent cell pairing events as measured by lattice light-sheet microscopy. **B.** Quantitative analysis of the nuclear N2-C concentration (nM) before sender cell contact (0 min), and at 30 and 60 min after contact. Lines connect data points from the same cell pairing event (n = 9, same nuclei from **A**).

**Figure S11.**
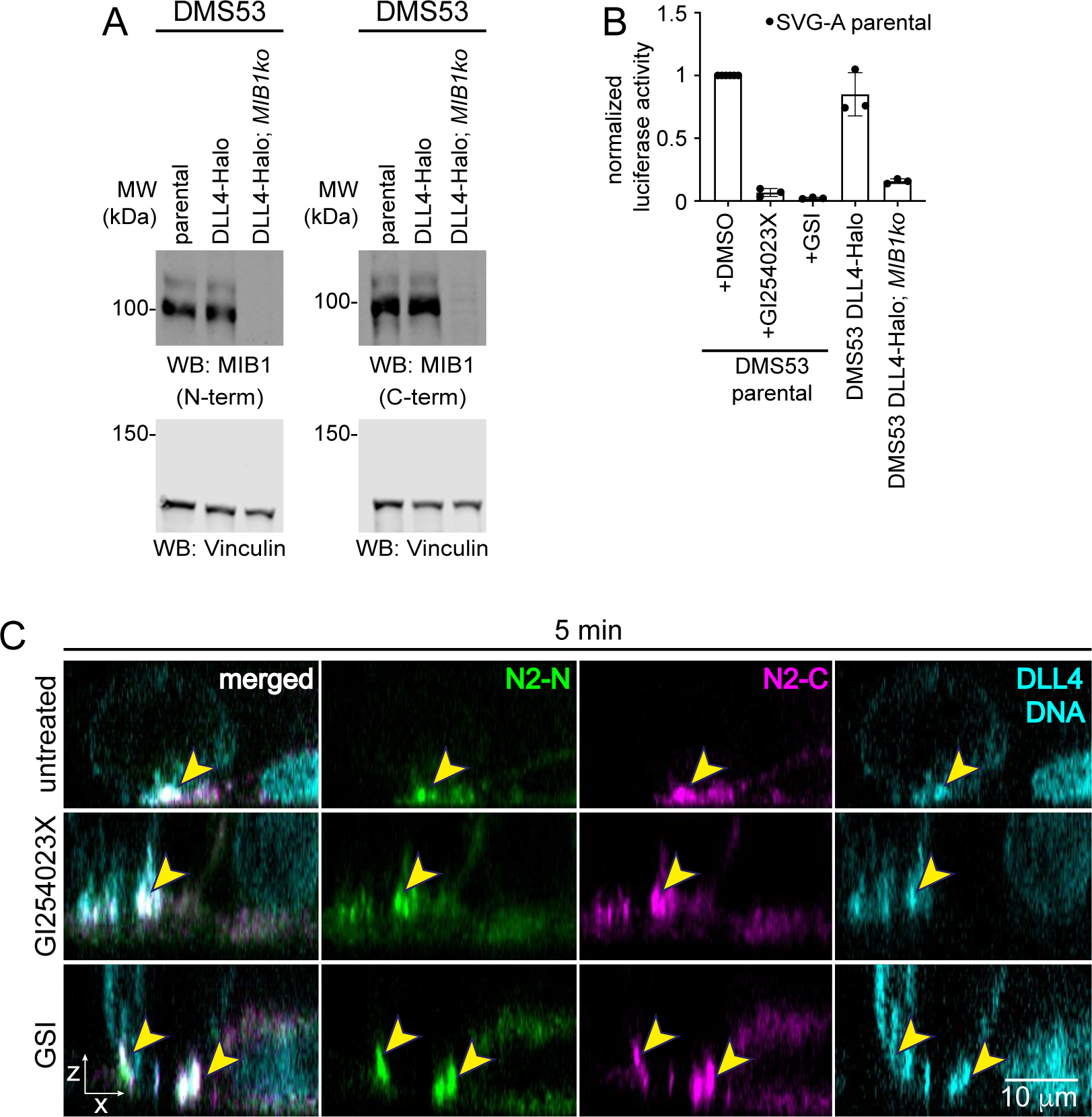
Characterization of DMS53 DLL4-HaloTag *MIB1ko* cells and influence of protease inhibitors on synapse formation. **A.** Western blots probing for MIB1 or vinculin in DMS53 parental cells, DMS53 DLL4-HaloTag cells, and DMS53 DLL4- HaloTag *MIB1ko* cells. MIB1 knockout was achieved by excision of the first exon of *MIB1*. Immunodetection of MIB1 was performed using two different antibodies recognizing N- or C-terminal regions of MIB1. Detection of Vinculin was used as a loading control. **B.** Luciferase reporter gene assay. Parental SVG-A cells were co-transfected with a reporter plasmid containing firefly luciferase under control of the Notch-responsive TP1 promoter (Kurooka et al., 1998; Minoguchi et al., 1997) and an internal control Renilla luciferase plasmid (Promega) using Lipofectamine™ 2000 (Invitrogen). 24 hours after transfection, these cells were co-cultured with DMS53 parental cells in the presence of DMSO, GI254023X (5 μM) or GSI (0.5 μM), with DMS53 DLL4-HaloTag cells, or with DMS53 DLL4-Halo;*MIB1ko* cells. Cells were lysed 24 hours later, and the firefly and Renilla luciferase activity of each lysate was measured using a Dual Luciferase Assay Kit (Promega). The firefly/Renilla ratio was normalized to the signal for co-culture of SVG-A parental cells with DMS53 cells in the presence of DMSO. Data are plotted as mean ± standard deviation, n = 3 independent biological replicates. **C**. Representative orthogonal view single plane images acquired using the spinning disk microscope of synapses between NOTCH2 and DLL4 cells when untreated, treated with GI254023X, or GSI. N2- N is green and N2-C is magenta. DLL4 and the nucleus of the NOTCH2 cell are in cyan. Yellow arrows point to the synapses in the merged and single channel images. Scale bar as indicated.

## References

Agrawal, N., Frederick, M. J., Pickering, C. R., Bettegowda, C., Chang, K., Li, R. J., Fakhry, C., Xie, T.-X., Zhang, J., Wang, J., Zhang, N., El-Naggar, A. K., Jasser, S. A., Weinstein, J. N., Treviño, L., Drummond, J. A., Muzny, D. M., Wu, Y., Wood, L. D., … Myers, J. N. (2011). Exome sequencing of head and neck squamous cell carcinoma reveals inactivating mutations in NOTCH1. Science, 333(6046), 1154– 1157. 10.1126/science.1206923

Aguet, F., Upadhyayula, S., Gaudin, R., Chou, Y. Y., Cocucci, E., He, K., Chen, B. C., Mosaliganti, K., Pasham, M., Skillern, W., Legant, W. R., Liu, T. L., Findlay, G., Marino, E., Danuser, G., Megason, S., Betzig, E., & Kirchhausen, T. (2016). Membrane dynamics of dividing cells imaged by lattice light-sheet microscopy. Molecular Biology of the Cell, 27(22), 3418–3435. 10.1091/mbc.E16-03-0164

Allgood, A. G., & Barrick, D. (2011). Mapping the Deltex-binding surface on the notch ankyrin domain using analytical ultracentrifugation. Journal of Molecular Biology, 414(2), 243–259. 10.1016/j.jmb.2011.09.050

Arnett, K. L., Hass, M., McArthur, D. G., Ilagan, M. X. G., Aster, J. C., Kopan, R., & Blacklow, S. C. (2010). Structural and mechanistic insights into cooperative assembly of dimeric Notch transcription complexes. Nature Structural & Molecular Biology, 17(11), 1312–1317. 10.1038/nsmb.1938

Artavanis-Tsakonas, S., Muskavitcht, M. A. T., & Yedvobnick, B. (1983). Molecular cloning of Notch, a locus affecting neurogenesis in Drosophila melanogaster. Genetics, 80, 1977–1981.

Aster, J. C., Pear, W. S., & Blacklow, S. C. (2017). The varied roles of Notch in cancer. Annual Review of Pathology, 12, 245–275. 10.1146/annurev-pathol-052016-100127

Blaumueller, C. M., Qi, H., Zagouras, P., & Artavanis-Tsakonas, S. (1997). Intracellular cleavage of Notch leads to a heterodimeric receptor on the plasma membrane. Cell, 90(2), 281–291. 10.1016/s0092-8674(00)80336-0

Bray, S. J. (2016). Notch signalling in context. Nature Reviews. Molecular Cell Biology, 17(11), 722–735. 10.1038/nrm.2016.94

Chapman, G., Major, J. A., Iyer, K., James, A. C., Pursglove, S. E., Moreau, J. L. M., & Dunwoodie, S. L. (2016). Notch1 endocytosis is induced by ligand and is required for signal transduction. Biochimica et Biophysica Acta, 1863(1), 166–177. 10.1016/j.bbamcr.2015.10.021

Chapman, G., Sparrow, D. B., Kremmer, E., & Dunwoodie, S. L. (2011). Notch inhibition by the ligand DELTA-LIKE 3 defines the mechanism of abnormal vertebral segmentation in spondylocostal dysostosis. Human Molecular Genetics, 20(5), 905–916. 10.1093/hmg/ddq529

Chastagner, P., Rubinstein, E., & Brou, C. (2017). Ligand-activated Notch undergoes DTX4-mediated ubiquitylation and bilateral endocytosis before ADAM10 processing. Science Signaling, 10(483), 1–14. 10.1126/scisignal.aag2989

Chen, B.-C., Legant, W. R., Wang, K., Shao, L., Milkie, D. E., Davidson, M. W., Janetopoulos, C., Wu, X. S., Hammerv, J. A., Iii, Liu, Z., English, B. P., Mimori-Kiyosue, Y., Romero, D. P., Ritter, A. T., Lippincott-Schwartz, J., Fritz-Laylin, L., Mullins, R. D., Mitchell, D. M., Bembenek, J. N., … Betzig, E. (2014). Lattice Light Sheet Microscopy: Imaging Molecules to Embryos at High Spatiotemporal Resolution Bi-Chang. Science, 346(6208). 10.1126/science.1257998.Lattice

Chen, E. Y., Tan, C. M., Kou, Y., Duan, Q., Wang, Z., Meirelles, G. V., Clark, N. R., & Ma’ayan, A. (2013). Enrichr: interactive and collaborative HTML5 gene list enrichment analysis tool. BMC Bioinformatics, 14, 128. 10.1186/1471-2105-14-128

Chou, Y.-Y., Krupp, A., Kaynor, C., Gaudin, R., Ma, M., Cahir-McFarland, E., & Kirchhausen, T. (2016). Inhibition of JCPyV infection mediated by targeted viral genome editing using CRISPR/Cas9. Scientific Reports, 6, 36921. 10.1038/srep36921

Couturier, L., Vodovar, N., & Schweisguth, F. (2012). Endocytosis by Numb breaks Notch symmetry at cytokinesis. Nature Cell Biology, 14(2), 131–139. 10.1038/ncb2419

Daskalaki, A., Shalaby, N. A., Kux, K., Tsoumpekos, G., Tsibidis, G. D., Muskavitch, M. A. T., & Delidakis, C. (2011). Distinct intracellular motifs of Delta mediate its ubiquitylation and activation by Mindbomb1 and Neuralized. The Journal of Cell Biology, 195(6), 1017–1031. 10.1083/jcb.201105166

Dobin, A., Davis, C. A., Schlesinger, F., Drenkow, J., Zaleski, C., Jha, S., Batut, P., Chaisson, M., & Gingeras, T. R. (2012). STAR: ultrafast universal RNA-seq aligner. Bioinformatics, 29(1), 15–21. 10.1093/bioinformatics/bts635

Edgar, R., Domrachev, M., & Lash, A. E. (2002). Gene Expression Omnibus: NCBI gene expression and hybridization array data repository. Nucleic Acids Research, 30(1), 207–210. 10.1093/nar/30.1.207

Falo-Sanjuan, J., & Bray, S. (2022). Notch-dependent and -independent transcription are modulated by tissue movements at gastrulation. In eLife (Vol. 11). 10.7554/elife.73656

Falo-Sanjuan, J., Lammers, N. C., Garcia, H. G., & Bray, S. J. (2019). Enhancer Priming Enables Fast and Sustained Transcriptional Responses to Notch Signaling. Developmental Cell, 50(4), 411–425.e8. 10.1016/j.devcel.2019.07.002

Fehon, R. G., Kooh, P. J., Rebay, I., Regan, C. L., Xu, T., Muskavitch, M. A., & Artavanis-Tsakonas, S. (1990). Molecular interactions between the protein products of the neurogenic loci Notch and Delta, two EGF-homologous genes in Drosophila. Cell, 61(3), 523–534. 10.1016/0092-8674(90)90534-l

Gordon, W. R., Roy, M., Vardar-Ulu, D., Garfinkel, M., Mansour, M. R., Aster, J. C., & Blacklow, S. C. (2009). Structure of the Notch1-negative regulatory region: Implications for normal activation and pathogenic signaling in T-ALL. Blood, 113(18), 4381–4390. 10.1182/blood-2008-08-174748

Gordon, W. R., Vardar-Ulu, D., Histen, G., Sanchez-Irizarry, C., Aster, J. C., & Blacklow, S. C. (2007). Structural basis for autoinhibition of Notch. Nature Structural & Molecular Biology, 14(4), 295–300. 10.1038/nsmb1227

Govindaraj, K., & Post, J. N. (2021). Using FRAP to Quantify Changes in Transcription Factor Dynamics After Cell Stimulation: Cell Culture, FRAP, Data Analysis, and Visualization. Methods in Molecular Biology, 2221, 109–139. 10.1007/978-1-0716-0989-7_9

Grimm, J. B., Xie, L., Casler, J. C., Patel, R., Tkachuk, A. N., Choi, H., Lippincott-Schwartz, J., Brown, T. A., Glick, B. S., Liu, Z., & Lavis, L. D. (2020). Deuteration improves small-molecule fluorophores. In bioRxiv (p. 2020.08.17.250027). 10.1101/2020.08.17.250027

Guo, B., McMillan, B. J., & Blacklow, S. C. (2016). Structure and function of the Mind bomb E3 ligase in the context of Notch signal transduction. Current Opinion in Structural Biology, 41, 38–45. 10.1016/j.sbi.2016.05.012

Ilagan, M. X. G., Lim, S., Fulbright, M., Piwnica-Worms, D., & Kopan, R. (2011). Real-time imaging of Notch activation with a luciferase complementation-based reporter. Science Signaling, 4(181), 1–15. 10.1126/scisignal.2001656

Jacobson, K., Ishihara, A., & Inman, R. (1987). Lateral diffusion of proteins in membranes. Annual Review of Physiology, 49, 163–175. 10.1146/annurev.ph.49.030187.001115

Joutel, A., Corpechot, C., Ducros, A., Vahedi, K., Chabriat, H., Mouton, P., Alamowitch, S., Domenga, V., Cécillion, M., Marechal, E., Maciazek, J., Vayssiere, C., Cruaud, C., Cabanis, E. A., Ruchoux, M. M., Weissenbach, J., Bach, J. F., Bousser, M. G., & Tournier-Lasserve, E. (1996). Notch3 mutations in CADASIL, a hereditary adult-onset condition causing stroke and dementia. Nature, 383(6602), 707–710. 10.1038/383707a0

Kamath, B. M., Bauer, R. C., Loomes, K. M., Chao, G., Gerfen, J., Hutchinson, A., Hardikar, W., Hirschfield, G., Jara, P., Krantz, I. D., Lapunzina, P., Leonard, L., Ling, S., Ng, V. L., Hoang, P. L., Piccoli, D. A., & Spinner, N. B. (2012). NOTCH2 mutations in Alagille syndrome. Journal of Medical Genetics, 49(2), 138–144. 10.1136/jmedgenet-2011-100544

Kang, M., Day, C. A., Kenworthy, A. K., & DiBenedetto, E. (2012). Simplified equation to extract diffusion coefficients from confocal FRAP data. Traffic, 13(12), 1589–1600. 10.1111/tra.12008

Khait, I., Orsher, Y., Golan, O., Binshtok, U., Gordon-Bar, N., Amir-Zilberstein, L., & Sprinzak, D. (2016). Quantitative Analysis of Delta-like 1 Membrane Dynamics Elucidates the Role of Contact Geometry on Notch Signaling. Cell Reports, 14(2), 225–233. 10.1016/j.celrep.2015.12.040

Khamaisi, B., Luca, V. C., Blacklow, S. C., & Sprinzak, D. (2022). Functional Comparison between Endogenous and Synthetic Notch Systems. ACS Synthetic Biology, 11(10), 3343–3353. 10.1021/acssynbio.2c00247

Klueg, K. M., & Muskavitch, M. a. (1999). Ligand-receptor interactions and trans-endocytosis of Delta, Serrate and Notch: members of the Notch signalling pathway in Drosophila. Journal of Cell Science, 112 ( *Pt 19*), 3289–3297. 10.1038/ncb0609-678

Kovall, R. A., Gebelein, B., Sprinzak, D., & Kopan, R. (2017). The Canonical Notch Signaling Pathway: Structural and Biochemical Insights into Shape, Sugar, and Force. Developmental Cell, 41(3), 228–241. 10.1016/j.devcel.2017.04.001

Kuleshov, M. V., Jones, M. R., Rouillard, A. D., Fernandez, N. F., Duan, Q., Wang, Z., Koplev, S., Jenkins, S. L., Jagodnik, K. M., Lachmann, A., McDermott, M. G., Monteiro, C. D., Gundersen, G. W., & Ma’ayan, A. (2016). Enrichr: a comprehensive gene set enrichment analysis web server 2016 update. Nucleic Acids Research, 44(W1), W90–7. 10.1093/nar/gkw377

Kurooka, H., Kuroda, K., & Honjo, T. (1998). Roles of the ankyrin repeats and C-terminal region of the mouse Notch1 intracellular region. Nucleic Acids Research, 26(23), 5448–5455. 10.1093/nar/26.23.5448

Langmead, B., Trapnell, C., Pop, M., & Salzberg, S. L. (2009). Ultrafast and memory-efficient alignment of short DNA sequences to the human genome. Genome Biology, 10(3), R25. 10.1186/gb-2009-10-3-r25

Langridge, P. D., & Struhl, G. (2017). Epsin-Dependent Ligand Endocytosis Activates Notch by Force. Cell, 171(6), 1383–1396.e12. 10.1016/j.cell.2017.10.048

Li, L., Krantz, I. D., Deng, Y., Genin, A., Banta, A. B., Collins, C. C., Qi, M., Trask, B. J., Kuo, W. L., Cochran, J., Costa, T., Pierpont, M. E., Rand, E. B., Piccoli, D. A., Hood, L., & Spinner, N. B. (1997). Alagille syndrome is caused by mutations in human Jagged1, which encodes a ligand for Notch1. Nature Genetics, 16(3), 243–251. 10.1038/ng0797-243

Liao, Y., Smyth, G. K., & Shi, W. (2013). featureCounts: an efficient general purpose program for assigning sequence reads to genomic features. Bioinformatics, 30(7), 923–930. 10.1093/bioinformatics/btt656

Logeat, F., Bessia, C., Brou, C., LeBail, O., Jarriault, S., Seidah, N. G., & Israël, A. (1998). The Notch1 receptor is cleaved constitutively by a furin-like convertase. Proceedings of the National Academy of Sciences of the United States of America, 95(14), 8108–8112. 10.1073/pnas.95.14.8108

Los, G. V., Encell, L. P., McDougall, M. G., Hartzell, D. D., Karassina, N., Zimprich, C., Wood, M. G., Learish, R., Ohana, R. F., Urh, M., Simpson, D., Mendez, J., Zimmerman, K., Otto, P., Vidugiris, G., Zhu, J., Darzins, A., Klaubert, D. H., Bulleit, R. F., & Wood, K. V. (2008). HaloTag: a novel protein labeling technology for cell imaging and protein analysis. ACS Chemical Biology, 3(6), 373–382. 10.1021/cb800025k

Love, M. I., Huber, W., & Anders, S. (2014). Moderated estimation of fold change and dispersion for RNA-seq data with DESeq2. Genome Biology, 15(12), 550. 10.1186/s13059-014-0550-8

Malecki, M. J., Sanchez-Irizarry, C., Mitchell, J. L., Histen, G., Xu, M. L., Aster, J. C., & Blacklow, S. C. (2006). Leukemia-associated mutations within the NOTCH1 heterodimerization domain fall into at least two distinct mechanistic classes. Molecular and Cellular Biology, 26(12), 4642–4651. 10.1128/MCB.01655-05

Martin, A. P., Bradshaw, G. A., Eisert, R. J., Egan, E. D., Tveriakhina, L., Rogers, J. M., Dates, A. N., Scanavachi, G., Aster, J. C., Kirchhausen, T., Kalocsay, M., & Blacklow, S. C. (2023). A spatiotemporal Notch interaction map from plasma membrane to nucleus. Science Signaling, 16(796), eadg6474. 10.1126/scisignal.adg6474

Martincorena, I., Roshan, A., Gerstung, M., Ellis, P., Van Loo, P., McLaren, S., Wedge, D. C., Fullam, A., Alexandrov, L. B., Tubio, J. M., Stebbings, L., Menzies, A., Widaa, S., Stratton, M. R., Jones, P. H., & Campbell, P. J. (2015). Tumor evolution. High burden and pervasive positive selection of somatic mutations in normal human skin. Science, 348(6237), 880–886. 10.1126/science.aaa6806

McMillan, B. J., Schnute, B., Ohlenhard, N., Zimmerman, B., Miles, L., Beglova, N., Klein, T., & Blacklow, S. C. (2015). A Tail of Two Sites: A Bipartite Mechanism for Recognition of Notch Ligands by Mind Bomb E3 Ligases. Molecular Cell, 57(5), 912–924. 10.1016/j.molcel.2015.01.019

Meloty-Kapella, L., Shergill, B., Kuon, J., Botvinick, E., & Weinmaster, G. (2012). Notch ligand endocytosis generates mechanical pulling force dependent on dynamin, epsins, and actin. Developmental Cell, 22(6), 1299–1312. 10.1016/j.devcel.2012.04.005

Minoguchi, S., Taniguchi, Y., Kato, H., Okazaki, T., Strobl, L. J., Zimber-Strobl, U., Bornkamm, G. W., & Honjo, T. (1997). RBP-L, a transcription factor related to RBP- Jkappa. Molecular and Cellular Biology, 17(5), 2679–2687. 10.1128/MCB.17.5.2679

Mumm, J. S., Schroeter, E. H., Saxena, M. T., Griesemer, A., Tian, X., Pan, D. J., Ray, W. J., & Kopan, R. (2000). A ligand-induced extracellular cleavage regulates gamma-secretase-like proteolytic activation of Notch1. Molecular Cell, 5(2), 197–206. 10.1016/S1097-2765(00)80416-5

Nichols, J. T., Miyamoto, A., Olsen, S. L., D’Souza, B., Yao, C., & Weinmaster, G. (2007). DSL ligand endocytosis physically dissociates Notch1 heterodimers before activating proteolysis can occur. The Journal of Cell Biology, 176(4), 445–458. 10.1083/jcb.200609014

Oda, T., Elkahloun, A. G., Pike, B. L., Okajima, K., Krantz, I. D., Genin, A., Piccoli, D. A., Meltzer, P. S., Spinner, N. B., Collins, F. S., & Chandrasekharappa, S. C. (1997). Mutations in the human Jagged1 gene are responsible for Alagille syndrome. Nature Genetics, 16(3), 235–242. 10.1038/ng0797-235

Okano, M., Matsuo, H., Nishimura, Y., Hozumi, K., Yoshioka, S., Tonoki, A., & Itoh, M. (2016). Mib1 modulates dynamin 2 recruitment via Snx18 to promote Dll1 endocytosis for efficient Notch signaling. Genes to Cells: Devoted to Molecular & Cellular Mechanisms, 21(5), 425–441. 10.1111/gtc.12350

Parks, A. L., Klueg, K. M., Stout, J. R., & Muskavitch, M. A. (2000). Ligand endocytosis drives receptor dissociation and activation in the Notch pathway. Development, 127(7), 1373–1385. 10.1242/dev.127.7.1373

Pillidge, Z., & Bray, S. J. (2019). SWI/SNF chromatin remodeling controls Notch-responsive enhancer accessibility. EMBO Reports, 20(5), e46944. 10.15252/embr.201846944

Puente, X. S., Pinyol, M., Quesada, V., Conde, L., Ordóñez, G. R., Villamor, N., Escaramis, G., Jares, P., Beà, S., González-Díaz, M., Bassaganyas, L., Baumann, T., Juan, M., López-Guerra, M., Colomer, D., Tubío, J. M. C., López, C., Navarro, A., Tornador, C., … Campo, E. (2011). Whole-genome sequencing identifies recurrent mutations in chronic lymphocytic leukaemia. Nature, 475(7354), 101–105. 10.1038/nature10113

Salman, M. M., Marsh, G., Kusters, I., Delincé, M., Di Caprio, G., Upadhyayula, S., de Nola, G., Hunt, R., Ohashi, K. G., Gray, T., Shimizu, F., Sano, Y., Kanda, T., Obermeier, B., & Kirchhausen, T. (2020). Design and Validation of a Human Brain Endothelial Microvessel-on-a-Chip Open Microfluidic Model Enabling Advanced Optical Imaging. Frontiers in Bioengineering and Biotechnology, 8, 573775. 10.3389/fbioe.2020.573775

Schindelin, J., Arganda-Carreras, I., Frise, E., Kaynig, V., Longair, M., Pietzsch, T., Preibisch, S., Rueden, C., Saalfeld, S., Schmid, B., Tinevez, J.-Y., White, D. J., Hartenstein, V., Eliceiri, K., Tomancak, P., & Cardona, A. (2012). Fiji: an open-source platform for biological-image analysis. *Nature Methods*, *9*(7), 676–682. 10.1038/nmeth.2019

Shaner, N. C., Lambert, G. G., Chammas, A., Ni, Y., Cranfill, P. J., Baird, M. A., Sell, B. R., Allen, J. R., Day, R. N., Israelsson, M., Davidson, M. W., & Wang, J. (2013). A bright monomeric green fluorescent protein derived from Branchiostoma lanceolatum. Nature Methods, 10(5), 407–409. 10.1038/nmeth.2413

Sprinzak, D., & Blacklow, S. C. (2021). Biophysics of Notch Signaling. Annual Review of Biophysics. 10.1146/annurev-biophys-101920-082204

van Tetering, G., van Diest, P., Verlaan, I., van der Wall, E., Kopan, R., & Vooijs, M. (2009). Metalloprotease ADAM10 is required for Notch1 site 2 cleavage. The Journal of Biological Chemistry, 284(45), 31018–31027. 10.1074/jbc.M109.006775

Wang, N. J., Sanborn, Z., Arnett, K. L., Bayston, L. J., Liao, W., Proby, C. M., Leigh, I. M., Collisson, E. A., Gordon, P. B., Jakkula, L., Pennypacker, S., Zou, Y., Sharma, M., North, J. P., Vemula, S. S., Mauro, T. M., Neuhaus, I. M., Leboit, P. E., Hur, J. S., … Cho, R. J. (2011). Loss-of-function mutations in Notch receptors in cutaneous and lung squamous cell carcinoma. Proceedings of the National Academy of Sciences of the United States of America, 108(43), 17761–17766. 10.1073/pnas.1114669108

Weng, A. P., Ferrando, A. A., Lee, W., Morris, J. P., 4th, Silverman, L. B., Sanchez-Irizarry, C., Blacklow, S. C., Look, A. T., & Aster, J. C. (2004). Activating mutations of NOTCH1 in human T cell acute lymphoblastic leukemia. Science, 306(5694), 269–271. 10.1126/science.1102160

Wilhelm, J., Kühn, S., Tarnawski, M., Gotthard, G., Tünnermann, J., Tänzer, T., Karpenko, J., Mertes, N., Xue, L., Uhrig, U., Reinstein, J., Hiblot, J., & Johnsson, K. (2021). Kinetic and Structural Characterization of the Self-Labeling Protein Tags HaloTag7, SNAP-tag, and CLIP-tag. Biochemistry, 60(33), 2560–2575. 10.1021/acs.biochem.1c00258

Xu, X., Choi, S. H., Hu, T., Tiyanont, K., Habets, R., Groot, A. J., Vooijs, M., Aster, J. C., Chopra, R., Fryer, C., & Blacklow, S. C. (2015). Insights into Autoregulation of Notch3 from Structural and Functional Studies of Its Negative Regulatory Region. Structure, 23(7), 1227–1235. 10.1016/j.str.2015.05.001

